# Chromatin remodeler CHD4 establishes chromatin states required for ovarian reserve formation, maintenance, and germ cell survival

**DOI:** 10.1101/2024.08.12.607691

**Authors:** Yasuhisa Munakata, Mengwen Hu, Yuka Kitamura, Adam L. Bynder, Amelia S. Fritz, Richard M. Schultz, Satoshi H. Namekawa

## Abstract

The ovarian reserve defines female reproductive lifespan, which in humans spans decades due to the maintenance of meiotic arrest in non-growing oocytes (NGO) residing in primordial follicles. Unknown is how the chromatin state of NGOs is established to enable long-term maintenance of the ovarian reserve. Here, we show that a chromatin remodeler, CHD4, a member of the Nucleosome Remodeling and Deacetylase (NuRD) complex, establishes chromatin states required for formation and maintenance of the ovarian reserve. Conditional loss of CHD4 in perinatal mouse oocytes results in acute death of NGOs and depletion of the ovarian reserve. CHD4 establishes closed chromatin at regulatory elements of pro-apoptotic genes to prevent cell death and at specific genes required for meiotic prophase I to facilitate the transition from meiotic prophase I oocytes to meiotic arrested NGOs. In addition, CHD4 establishes closed chromatin at the regulatory elements of pro-apoptotic genes in male germ cells, allowing male germ cell survival. These results demonstrate a role for CHD4 in defining a chromatin state that ensures germ cell survival, thereby enabling the long-term maintenance of both female and male germ cells.

## Introduction

Germ cell maintenance and survival are fundamental for the continuous supply of gametes for reproduction. In adult mammals, while the male germline is maintained by self-renewal of spermatogonial stem cells, the female germline is not maintained by a stem cell-based mechanism but is maintained within a pool of meiotically arrested oocytes, called the ovarian reserve. These small non-growing oocytes (NGOs) residing in primordial follicles are arrested at the dictyate stage, the prolonged diplotene stage, of meiotic prophase I (MPI) ^1^. NGOs are the only source of fertilizable eggs throughout a female’s reproductive life span. Because the number of NGOs is finite, their premature depletion leads to infertility associated with early menopause, such as premature ovarian insufficiency (POI)^2^. However, the mechanisms underlying formation and maintenance of the ovarian reserve remain largely elusive.

In the female germline of mouse embryos, primordial germ cells (PGCs) initiate MPI after induction of genes specifically expressed in MPI (MPI genes)^3^. After completing chromosome synapsis and recombination, oocytes reach the dictyate stage around birth, gradually decreasing in number before formation of the ovarian reserve ^4, 5, 6^. During ovarian reserve formation, MPI genes are suppressed, and there is a transition in genome-wide transcription to become NGOs, which is termed the perinatal oocyte transition (POT)^7, 8^. POT is regulated by an epigenetic regulator, Polycomb Repressive Complex 1 (PRC1), to suppress MPI genes when oocytes exit MPI^7^. Concomitantly, an oocyte-specific transcription factor FIGLA and several signaling pathways, such as Notch, TGF-β, JNK, and hypoxia signaling that regulate gene expression are required for primordial follicle formation^9, 10, 11, 12, 13^. These studies raise the possibility that the chromatin state in NGOs is uniquely established to instruct the gene expression program for ovarian reserve formation.

To identify a chromatin-based mechanism underlying the process of the ovarian reserve formation and maintenance, we sought to examine the role of ATP-dependent chromatin remodelers that utilize the energy from ATP hydrolysis to reorganize chromatin and regulate gene expression^14, 15^. There are four major subfamilies of chromatin remodeling complexes, including SWI/SNF (switch/sucrose non-fermentable), ISWI (imitation SWI), NuRD (nucleosome remodeling and deacetylase)/CHD (chromodomain helicase DNA-binding)/mi-2, and INO80/SWR (SWI2/SNF2 related) families^14, 15^.

Among the ATPase subunits in these four major chromatin remodeler subfamilies, we focused our attention on CHD4 (also known as Mi-2β) based on its gene expression at POT and because CHD4 is associated with lineage commitment and differentiation processes^16, 17, 18, 19^. In the male germline, CHD4 regulates maintenance and survival of undifferentiated spermatogonia^20, 21, 22^. However, the molecular mechanisms underlying this process remain unknown.

Here we show that CHD4 has an essential function in formation and maintenance of the ovarian reserve and determine the molecular mechanisms common to both female and male germ cells. In the female germline, CHD4 establishes closed chromatin at regulatory elements of pro-apoptotic genes to prevent cell death and at MPI genes to facilitate the transition from MPI to NGO. Further, in the male germline, CHD4 establishes closed chromatin at the regulatory elements of pro-apoptotic genes, allowing male germ cell survival. Thus, CHD4 defines the chromatin state for maintenance of both female and male germ cells.

## Results

### CHD4 is required for ovarian reserve formation

To identify a key ATP-dependent chromatin remodeler that functions in ovarian reserve formation in mice, we compared gene expression profiles of ATPase subunits in representative chromatin remodeler subfamilies using previously published RNA-seq data^13^. Among these candidates, *Chd4* is highly expressed from embryonic day 14 (E14.5) oocytes in MPI to postnatal day 4 (P4) and P6 NGOs, which corresponds to the time of ovarian reserve formation when genome-wide gene expression changes occur during POT^7^ (Fig. 1a). *Chd4* expression was slightly downregulated when NGOs in primordial follicle (labeled as “small” in Fig. 1a) are activated to become growing oocytes (GOs) in primary follicles (labeled as “large” in Fig. 1a); this transition is termed the primordial to primary follicle transition (PPT)^13^, and the first wave occurs as early as P4 (Fig. 1a). Based on this gene expression profile, we focused on CHD4 and sought to determine the function of CHD4 in ovarian reserve formation.

**Fig 1.**
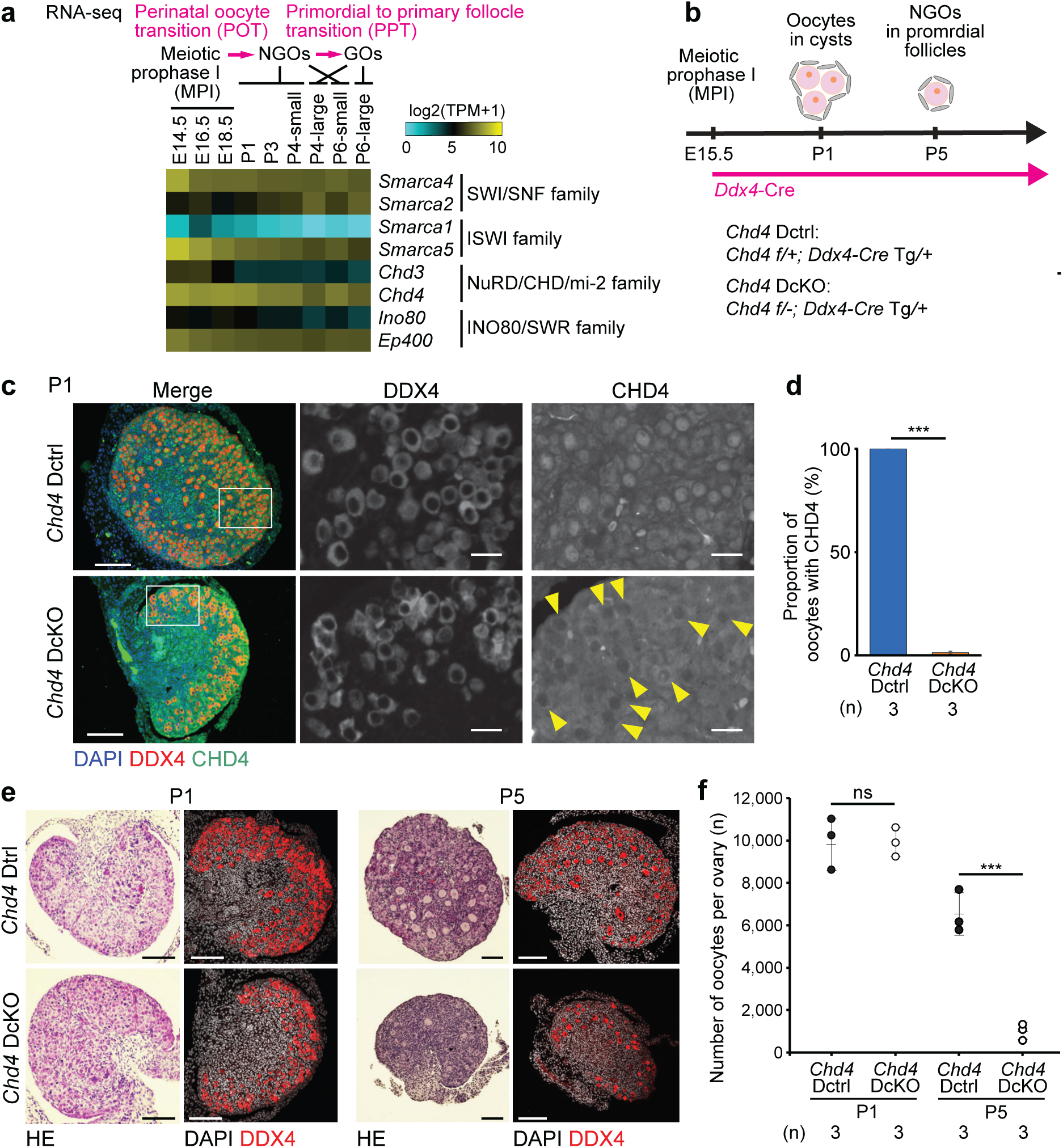
CHD4 deficiency causes oocyte loss. **a.** Heatmap showing bulk RNA-seq gene expression (log2 (TPM+1) values) for core subunits of ATP-dependent remodeling complexes in oocytes during oogenesis. Embryonic day (E) 18.5 to postnatal day (P) 3 indicate oocytes with meiosis in progress. P4 and P6 small indicate oocytes residing in primordial follicles. P4 and P6-large indicate growing oocytes in primary follicles. **b.** Schematic for mouse models and experiments. **c.** Immunostaining of DDX4 (red) and CHD4 (green) in ovaries of *Chd4* Dctrl and *Chd4* DcKO at P1. CHD4 is present only in somatic cells in *Chd4* DcKO. Bars: 100 μm (20 μm in the boxed area). Yellow arrowheads indicate oocytes with no CHD4 expression. **d.** Quantitative analysis of immunostaining. Percentages representing numbers of oocytes with CHD4 at P1. Data are presented as mean values ± SD. *** *P* < 0.001: Two-tailed unpaired t-tests. Three independent biological replicates were analyzed for each genotype. **e.** Ovarian sections of *Chd4* Dctrl and *Chd4* DcKO mice at P1 and P5, respectively. The sections were stained with hematoxylin and eosin or immunostained for DDX4 (red). Bars: 100 μm. Three mice were analyzed for each genotype at each time point, and representative images are shown. **f.** Dot plots showing the estimated numbers of oocytes per ovary from *Chd4* Dctrl and *Chd4* DcKO mice at P1 and P5, respectively. At least three mice were analyzed for each genotype at each time point. Central bars represent mean values. *** *P* < 0.001: ns, not significant; Two-tailed unpaired t-tests.

To determine the function of CHD4 in ovarian reserve formation, we generated *Chd4* conditional knockout mice using the *Chd4* floxed allele^23^ and *Ddx4*-Cre transgene, which is a germline-specific Cre line expressed from E15.5^24^ (*Chd4* ^f/-^; *Ddx4*-Cre ^Tg/+^; termed *Chd4* DcKO, Fig. 1b). CHD4 protein localized in the nucleus of the P1 oocytes in littermate controls (*Chd4* ^f/+^; *Ddx4*-Cre ^Tg/+^; termed *Chd4* Dctrl), but was absent in 98.8% of the P1 oocyte nuclei of *Chd4* DcKO mice (Fig1c, d), confirming the efficient deletion of CHD4 protein in *Chd4* DcKO oocytes. The estimated oocyte number of *Chd4* DcKO neonatal mice at P1 did not differ from that of *Chd4* Dctrl (Fig. 1e, f). However, by P5, when the ovarian reserve is established, the estimated oocyte number of *Chd4* DcKO newborn mice was markedly reduced compared to *Chd4* Dctrl (Fig. 1e, f). Apoptosis was likely responsible for the loss of oocytes in *Chd4* DcKO newborn mice because immunofluorescence staining for cleaved Caspase 3, a marker of apoptosis, revealed no difference in the proportion of cleaved Caspase 3-positive oocytes in P1, whereas the proportion of cleaved Caspase 3-positive oocytes in P3 *Chd4* DcKO ovaries was significantly increased compared to *Chd4* Dctrl (Fig. 2a, b). These results suggest that CHD4 is critical for ovarian reserve formation.

**Fig 2.**
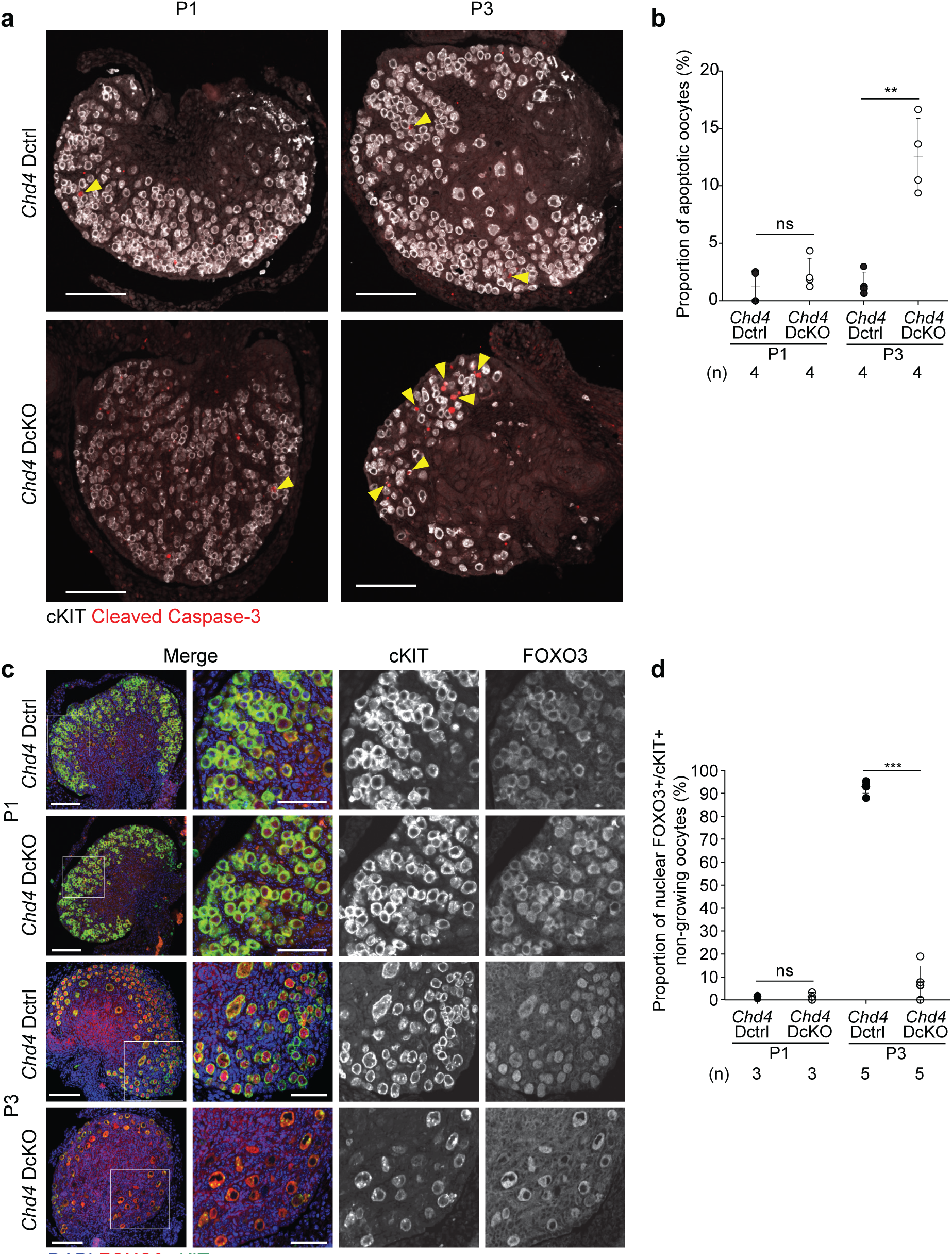
CHD4 is required for ovarian reserve formation. **a.** Immunofluorescence staining of cKIT (white) and Cleaved Caspase-3 (red) in ovaries of *Chd4* Dctrl and *Chd4* DcKO at P1 and P3, respectively. Yellow arrowheads indicate Cleaved Caspase-3^+^ apoptotic oocytes. Bars: 100 μm. Three mice were analyzed for each genotype at each time point, and representative images are shown. **b.** Quantitative analysis of immunostaining. Dot plots showing the percentages of Cleaved Caspase-3^+^ apoptotic oocytes per cKIT^+^ oocytes at P1 and P3. Four independent biological replicates were analyzed for each genotype at each time point. ** *P* < 0.01: ns, not significant; Two-tailed unpaired t-tests. **c.** Immunofluorescence staining of FOXO3 (red) and cKIT (green) in ovaries of *Chd4* Dctrl and *Chd4* DcKO at P1 and P5, respectively. Bars: 100 μm (50 μm in the boxed area). Three and five mice were analyzed for each genotype at each time point, and representative images are shown. **d.** Quantitative analysis of immunostaining. Dot plots showing the percentages of nuclear FOXO3^+^ oocytes per cKIT^+^ oocytes at P1 and P5, respectively. Three and five independent biological replicates were analyzed for each genotype at each time point. *** *P* < 0.001; ns, not significant; Two-tailed unpaired t-tests.

We next examined whether the ovarian reserve is properly established in *Chd4* DcKO ovaries. During the formation of NGOs in the ovarian reserve, localization of the transcription factor FOXO3 changes from the cytoplasm to the nucleu^25^; nuclear localization of FOXO3 is a hallmark of NGOs^25^.

Immunofluorescence staining for FOXO3 showed that in P1 FOXO3 was localized in the cytoplasm of most oocytes in both *Chd4* Dctrl and *Chd4* DcKO (Fig. 2c, d). However, at P3, FOXO3 was localized in the nucleus of 92.9% of oocytes in *Chd4* Dctrl whereas nuclear localization was only 7.9% of oocytes in *Chd4* DcKO had FOXO3 (Fig. 2c, d). Therefore, NGOs are not properly generated in *Chd4* DcKO ovaries. Nevertheless, the behavior of meiotic chromosomes in MPI, including progression to the dictyate stage of MPI, appeared normal in *Chd4* DcKO (Supplementary Fig. 1a). Thus, cell death is not initiated by defects in meiotic chromosome behavior.

### CHD4 represses MPI genes and apoptosis genes in ovarian reserve formation

To further examine the function of CHD4 in ovarian reserve formation, NGOs were isolated from P1 and P5 ovaries, and RNA-seq analysis was performed (Supplementary Fig. 1b). In P1 *Chd4* DcKO NGOs, 533 genes were up-regulated, and 348 genes were down-regulated (Fig. 3a, left, Supplementary Data 1). In P5 *Chd4* DcKO NGOs, 569 genes were up-regulated, and 396 genes were down-regulated (Fig. 3a, right, Supplementary Data 2). To infer possible functions of these differentially expressed genes, we performed Gene ontology enrichment analyses. The genes down-regulated in P1 and P5 *Chd4* DcKO oocytes were enriched with genes involved in “oogenesis” and “female gamete generation”. On the other hand, the genes up-regulated in P5 *Chd4* DcKO oocytes were enriched with genes involved in MPI, such as “homologous chromosome pairing at meiosis” (Fig. 3b), suggesting that CHD4 represses MPI genes. During ovarian reserve formation, MPI genes are repressed as oocytes exit from MPI and the fetal program^7^. Therefore, we examined how MPI genes are regulated in *Chd4* DcKO NGOs.

**Fig 3.**
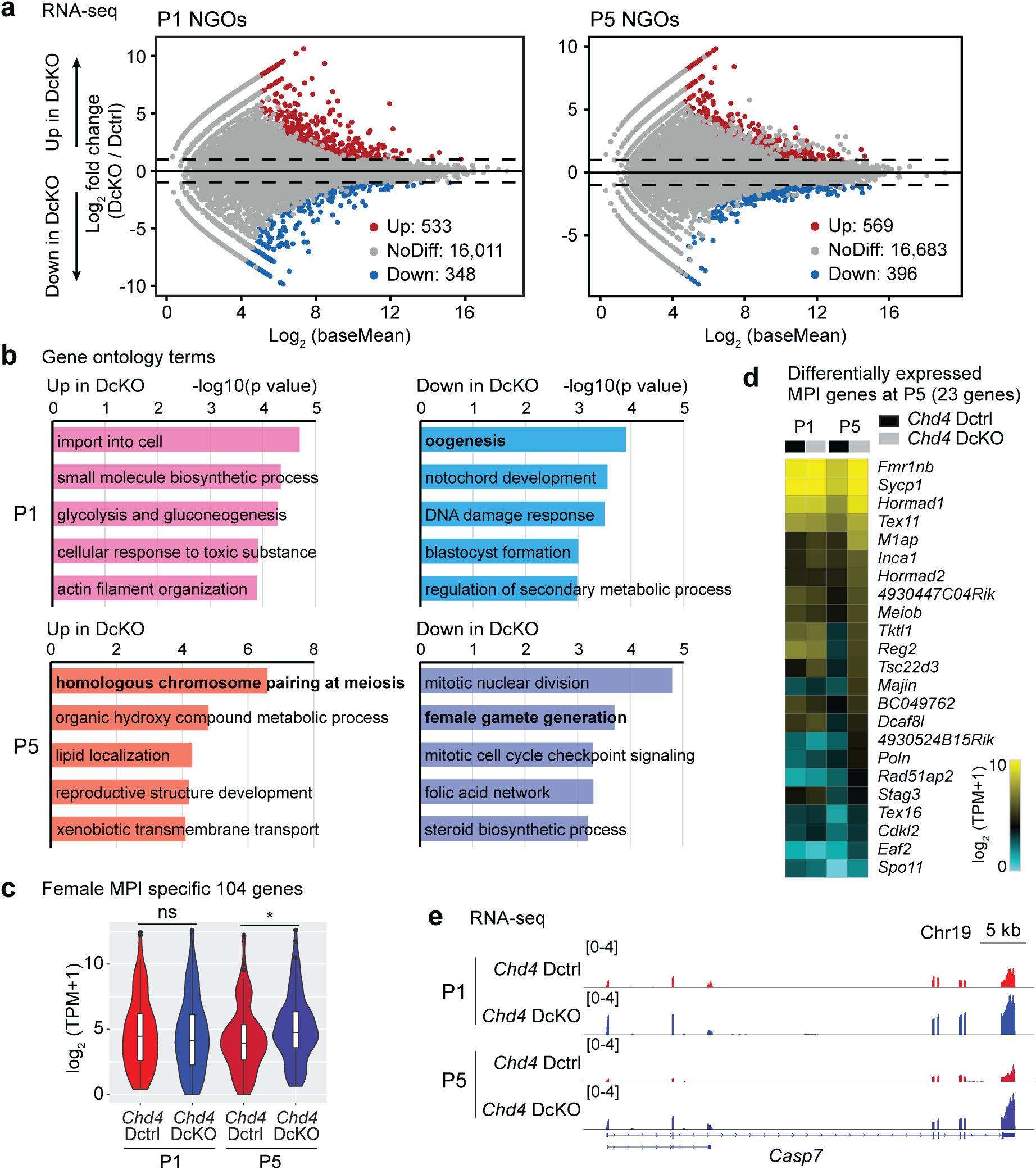
CHD4 represses meiotic prophase I genes and apoptosis-related genes. **a.** Comparison of transcriptomes between *Chd4* Dctrl and *Chd4* DcKO oocytes at P1 and P5, respectively. 500 non-growing oocytes (NGOs) isolated from P1 or P5 ovaries were pooled as one replicate, and two independent biological replicates were examined for RNA-seq. Differentially expressed genes (DEGs: Log2FoldChange > 1, Padj < 0.05, binominal test with Benjamini–Hochberg correction) are colored (red: upregulated in *Chd4* DcKO oocytes; blue: downregulated in *Chd4* DcKO oocytes). **b.** Gene ontology term enrichments analysis of differentially expressed genes detected in **a**. **c.** Violin plots with a Box plot indicate TPM values for female MPI-specific genes (104 genes) in *Chd4* Dctrl and *Chd4* DcKO oocytes at P1 and P5. The central lines represent medians. The upper and lower hinges correspond to the 25th and 75th percentiles. The upper and lower whiskers are extended from the hinge to the largest value no further than the 1.5x inter-quartile range (IQR) from the hinge. * *P* < 0.05: ns, not significant; Wilcoxon rank sum test. **d.** Heatmaps showing expression of the P5 up-regulated differentially expressed MPI-specific genes in *Chd4* DcKO oocytes at P1 and P5, respectively. **e.** RNA-seq track views at the *Casp7* gene locus. The y-axis represents normalized tag counts for bulk RNA-seq in each sample. Data ranges are shown in brackets.

In a previous study^26^, 104 genes were identified to be MPI-specific genes in fetal oocytes^26^. The expression of these MPI genes is comparable between *Chd4* DcKO and *Chd4* Dctrl NGOs at P1 (Fig. 3c). In contrast, in P5 NGOs, expression of MPI genes was significantly up-regulated in *Chd4* DcKO relative to *Chd4* Dctrl (Fig. 3c). Among them, 23 MPI genes, such as *Spo11*^27, 28^, *Sycp1*^29^, *Hormad1*^30, 31, 32^, *Meiob*^33^, and *Majin*^34^, which are important for MPI progression, were included in the differentially expressed genes in P5 NGOs (Fig. 3d). Because there is a genome-wide gene expression change in POT in normal oogenesis^7^, we next examined how differentially expressed genes at POT are regulated in the *Chd4* DcKO NGOs. In P5 *Chd4* DcKO NGOs, up-regulated genes at POT were down-regulated, while down-regulated genes at POT were up-regulated (Supplementary Fig. S1c, d, Supplementary Data 3), further confirming that POT is defective in accordance with defective ovarian reserve formation.

We also examined the expression of the genes in the mouse apoptosis pathway defined in the KEGG database^35^. Both in P1 and P5 NGOs, *Casp7*, a gene important for apoptosis, was significantly up-regulated (Fig. 3e). Taken together, CHD4 is required for the repression of MPI genes and apoptosis genes during ovarian reserve formation.

### CHD4 represses chromatin accessibility to down-regulate genes

Because CHD4 represses transcription and chromatin accessibility in various cell types^16, 36, 37^, we next sought to determine how CHD4 regulates chromatin accessibility during ovarian reserve formation. We used the assay for transposase-accessible chromatin by sequencing (ATAC-seq)^38, 39^ to assess the effect of CHD4 loss on chromatin accessibility in ovarian reserve formation (Supplementary Fig. 2a). A representative track view confirms that peak patterns are consistent between two biological replicates (Supplementary Fig. 2b). An ATAC-seq analysis of P1 NGOs showed that accessibility was massively increased in *Chd4* DcKO (Fig. 4a). These increased accessibility regions are mainly introns and intergenic regions, and a relatively minor change was observed at promoters and transcription termination sites (TTSs: Fig. 4b). This result suggests that CHD4 regulates distal cis-regulatory elements such as enhancers.

**Fig 4.**
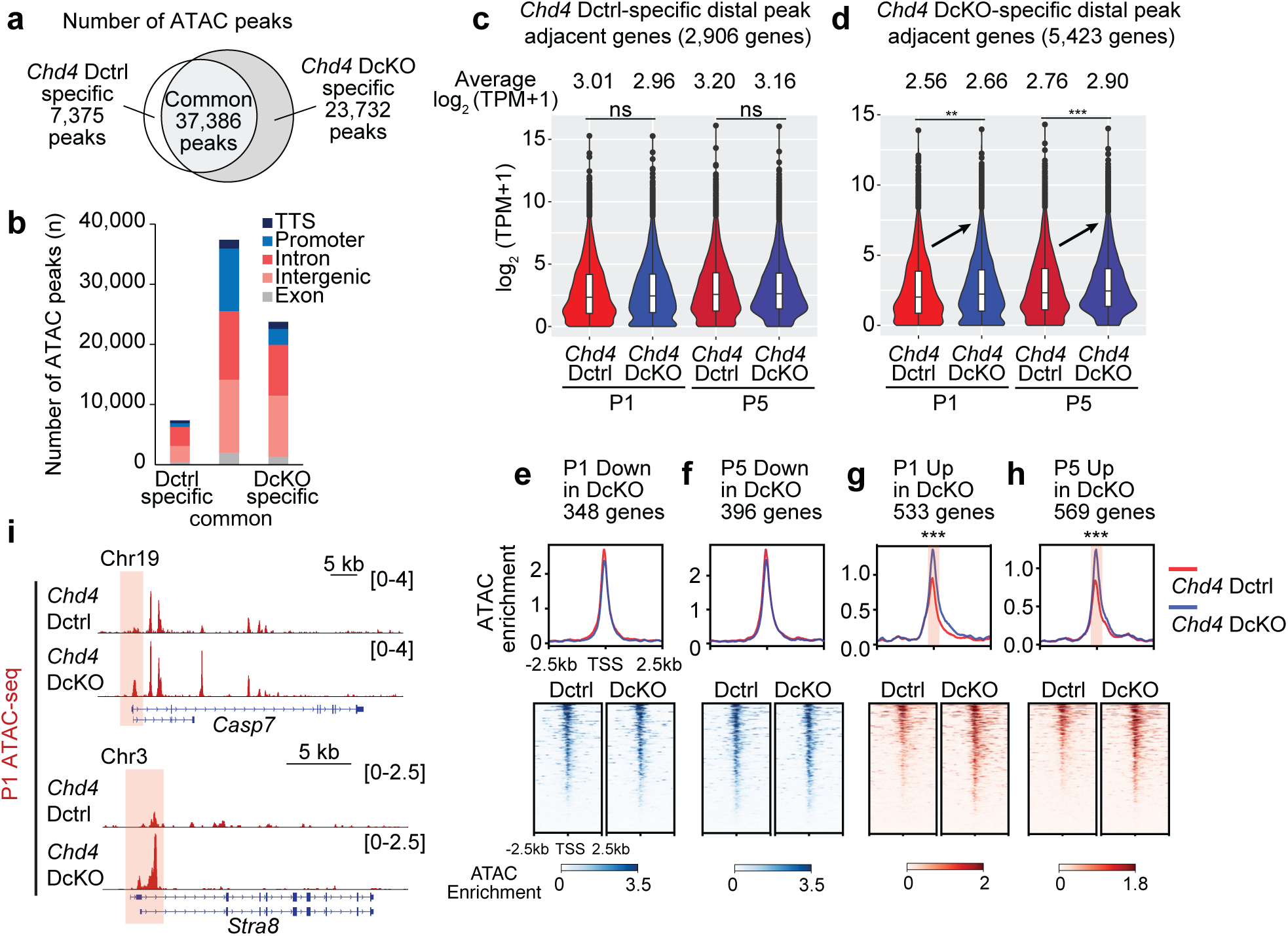
CHD4-dependent regulation of accessible chromatin in perinatal oocytes. **a.** Venn diagram indicates overlap of ATAC-seq peaks between *Chd4* Dctrl and *Chd4* DcKO oocytes at P1. **b.** Numbers and genomic distribution of ATAC-seq peaks in **a**. **c**, **d**. Violin plots with a box plot indicate changes in TPM values of genes adjacent to specific ATAC-seq peaks in *Chd4* Dctrl (**c**) and *Chd4* DcKO (**d**) oocytes at P1. The central lines represent medians. The upper and lower hinges correspond to the 25th and 75th percentiles. The upper and lower whiskera are extended from the hinge to the largest value no further than the 1.5x inter-quartile range (IQR) from the hinge. *** *P* < 0.001; ** *P* < 0.01; ns, not significant; Wilcoxon rank sum test. **e, f, g, h**. Heatmaps and average tag density plots of ATAC-seq enrichment around TSS (±2.5 kb) of downregulated in *Chd4* DcKO oocytes at P1 (**e**) and P5 (**f**) and upregulated in *Chd4* DcKO oocytes at P1 (**g**) and P5 (**h**). *** *P* < 0.001; Wilcoxon rank sum test. **i.** Representative track views of *Casp7* and *Stra8* gene loci show ATAC-seq signals in *Chd4* Dctrl and *Chd4* DcKO oocytes at P1. The y-axis represents normalized tag counts for ATAC-seq in each sample. The regions around TSSs are highlighted in red.

Next, we examined the relationship between changes in the distal accessible regions and changes in gene expression following CHD4 loss. Expression of 2,906 genes adjacent to *Chd4* Dctrl-specific distal accessible regions (outside the transcription start sites (TSSs) ±1 kb window) was similar between *Chd4* DcKO and *Chd4* Dctrl NGOs both in P1 and P5 (Fig. 4c). However, expression of 5,423 genes adjacent to *Chd4* DcKO-specific distal accessible regions was globally up-regulated in *Chd4* DcKO NGOs both in P1 and P5 (Fig. 4d). Therefore, CHD4 represses distal accessible regions to down-regulate genes. Further, we examined chromatin accessibility at the promoters of differentially expressed genes in *Chd4* DcKO NGOs. We found that up-regulated genes in *Chd4* DcKO NGOs are associated with increased accessibility at promoters both in P1 and P5, whereas down-regulated genes were not associated with changes in chromatin accessibility (Fig. 4e-h). Chromatin accessibility was increased at the TSS of genes whose expression was up-regulated in *Chd4* DcKO; for example, an apoptotic gene *Casp7*, and an MPI gene *Stra8*, which is the transcription factor critical for MPI gene expression^40^ (Fig. 4i). Together, we conclude that CHD4 represses chromatin accessibility both at promoters and distal regulatory elements to repress genes in ovarian reserve formation.

### CHD4 binds chromatin to regulate MPI and apoptosis genes

Because CHD4 deficiency causes a massive increase in chromatin accessibility, we hypothesize that CHD4 directly binds target sites to regulate chromatin accessibility. To test this hypothesis, we performed Cleavage Under Targets and Tagmentation (CUT&Tag) analysis^41^ on CHD4 using P1 oocytes to determine where CHD4 is bound in the genome (Supplementary Fig. 2c). CUT&Tag analysis revealed that the majority of CHD4 peaks were enriched in introns and intergenic regions (Fig. 5a). In addition, most of the CHD4 peaks were located 5-500 kb away from the TSSs (Fig. 5b), consistent with the genomic sites of accessibility changes in *Chd4* DcKO NGOs. We compared the ATAC peaks with the CHD4 peaks and found that, surprisingly, only a minor portion of them overlapped (Supplementary Fig. 2d), and this is the case for the ATAC distal peaks (Fig. 5c). We compared the adjacent gene expression near the CHD4 peaks and found that it was significantly increased in P5 *Chd4* DcKO (Fig. 5d), suggesting that CHD4 binds to repress target genes. However, counterintuitively, the CHD4 signals were enriched at the TSSs of the gene that was significantly down-regulated in P1 and P5 *Chd4* DcKO (Fig. 5e). These results suggest that CHD4 not only directly regulates chromatin accessibility but may also regulate gene expression without changing chromatin accessibility.

**Fig 5.**
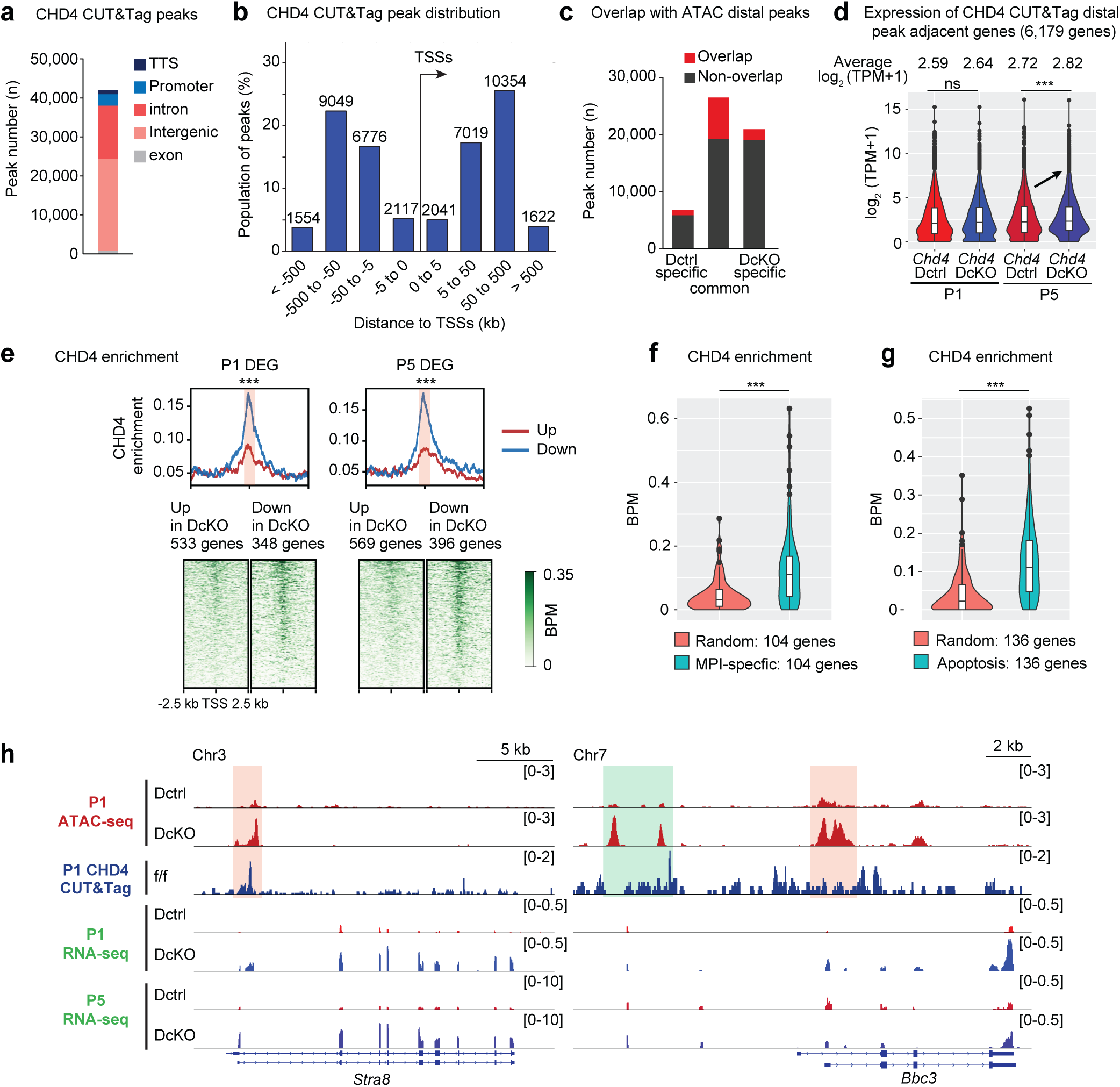
CHD4 binding sites in non-growing oocytes at P1. **a.** Numbers and genomic distribution of CHD4 CUT&Tag peaks in *Chd4* f/f oocytes at P1. **b.** Bar chart depicts the regional distribution of CHD4 CUT&Tag peaks to TSSs. **c.** Overlap between ATAC distal peaks (> 1kb from TSSs) and CHD4 CUT&Tag peaks within classified ATAC peaks. **d.** Violin plots with a box plot indicate changes in TPM values of genes adjacent to CHD4 CUT&Tag peaks in *Chd4* Dctrl and *Chd4* DcKO oocytes at P1 and P5. The central lines represent medians. The upper and lower hinges correspond to the 25th and 75th percentiles. The upper and lower whiskers are extended from the hinge to the largest value no further than the 1.5x inter-quartile range (IQR) from the hinge. *** *P* < 0.001; ns, not significant; Wilcoxon rank sum test. **e.** Heatmaps and average tag density plots of CHD4 enrichment around TSS (±2.5 kb) of DEG in *Chd4* Dctrl and *Chd4* DcKO oocytes at P1 and P5. *** *P* < 0.001; Wilcoxon rank sum test. **f, g**. Violin plots with a box plot indicate CHD4 enrichment around TSS (±1 kb) for female MPI-specific genes (**f**, 104 genes) and apoptosis pathway genes in the KEGG database (**g**, 136 genes) in *Chd4* Dctrl and *Chd4* DcKO oocytes at P1 and P5. The central lines represent medians. The upper and lower hinges correspond to the 25th and 75th percentiles. The upper and lower whiskers are extended from the hinge to the largest value no further than the 1.5x IQR from the hinge. *** *P* < 0.001; Wilcoxon rank sum test. **h**. Representative track views of *Stra8* and *Bbc3* loci in P1 and P5 oocytes of indicated genotypes. Data ranges are shown in brackets. Specific ATAC-seq peak regions in *Chd4* DcKO are highlighted.

To further elucidate the function of CHD4 in formation of ovarian reserve, we focused on apoptosis-associated genes and MPI genes whose expression was up-regulated in *Chd4* DcKO NGOs. CHD4 was enriched in the TSSs of the respective gene groups compared to randomly selected regions (Fig. 5f, g). At the *Stra8* gene locus, CHD4 binds the TSS, where chromatin accessibly increased in the *Chd4* DcKO NGOs (Fig. 5h, left). Furthermore, at the pro-apoptotic *Bbc3* (also known as *Puma*) gene locus, CHD4 bound not only at the TSS but also at the upstream region where chromatin accessibility was increased in *Chd4* DcKO NGOs (Fig. 5h, right). Thus, for some important target genes, CHD4 directly regulates chromatin accessibility, supporting a model in which CHD4 represses expression by regulating chromatin accessibility.

## CHD4 is required for the maintenance of the ovarian reserve and oocyte survival

Because a critical aspect of ovarian reserve is the long maintenance of chromatin states during the female reproductive life span, we next determined whether CHD4 is required for maintenance of ovarian reserve after its establishment. To elucidate the function of CHD4 in the maintenance of NGOs in the ovarian reserve, we generated another line of CHD4 conditional knockout mice using *Gdf9*-iCre, which is expressed NGOs from P3^42^ (*Chd4* ^f/f^; *Gdf9*-iCre ^Tg/+^; termed *Chd4* GcKO) (Fig. 6a). CHD4 was localized in the nuclei of NGOs in primordial follicles and GOs in primary follicles of P10 ovaries, and nearly complete depletion of CHD4 was observed in P10 *Chd4* GcKO oocytes (Supplementary Fig. 3a, b). In P10 ovaries in which primordial follicle formation is complete, the estimated numbers of NGOs and GOs in the ovaries of *Chd4* GcKO mice were significantly reduced in both NGOs and GOs compared to *Chd4* Gctrl mice (Fig. 6b, c). These results indicate that CHD4 is essential for the maintenance of NGOs and the survival of GOs.

**Fig 6.**
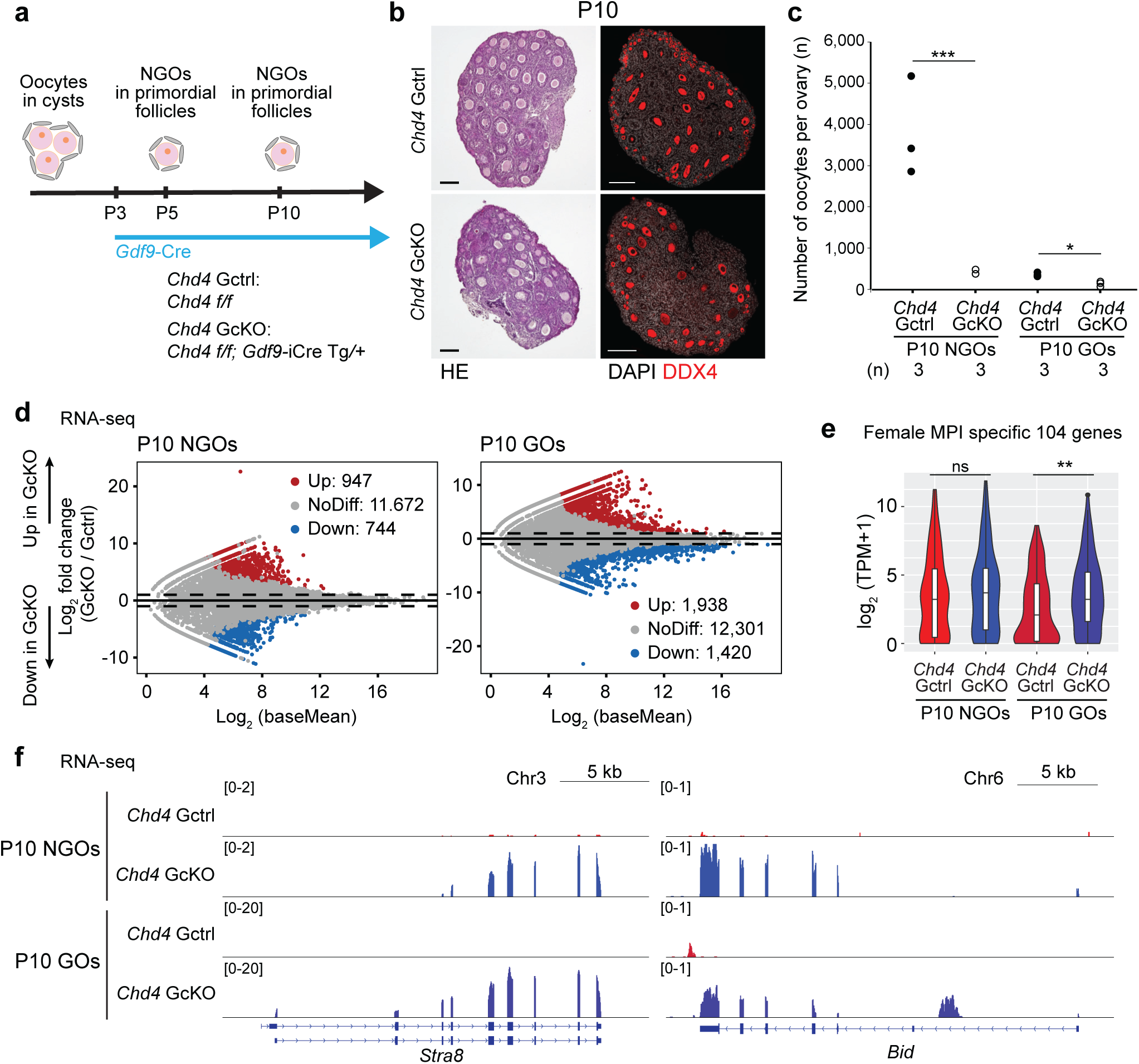
CHD4 is required for ovarian reserve maintenance. **a.** Schematic for mouse models and experiments. **b.** Ovarian sections of *Chd4* Gctrl and *Chd4* GcKO mice at P10. The sections were stained with hematoxylin and eosin or immunostained for DDX4 (red). Bars: 100 μm. Three mice were analyzed for each genotype at each time point, and representative images are shown. **c.** Dot plots showing the estimated numbers of oocytes per ovary from *Chd4* Gctrl and *Chd4* GcKO mice at P10. At least three mice were analyzed for each genotype at each time point. Central bars represent mean values. *** *P* < 0.001; * *P* < 0.05; Two-tailed unpaired t-tests. **d.** Comparison of transcriptomes between *Chd4* Gctrl and *Chd4* GcKO non-growing oocytes (NGOs) and growing oocytes (GOs) at P10. 500 NGOs and 100 GOs were isolated from P10 ovaries and were pooled as one replicate, and two independent biological replicates were examined for RNA-seq. Differentially expressed genes (DEGs: Log2FoldChange > 1, Padj < 0.05, binominal test with Benjamini–Hochberg correction) are colored (red: upregulated in *Chd4* GcKO oocytes; blue: downregulated in *Chd4* GcKO oocytes). **e.** Violin plots with a Box plot indicate TPM values for female MPI-specific genes (104 genes) in *Chd4* Gctrl and *Chd4* GcKO oocytes at P10. The central lines represent medians. The upper and lower hinges correspond to the 25th and 75th percentiles. The upper and lower whiskers are extended from the hinge to the largest value no further than the 1.5x inter-quartile range (IQR) from the hinge. ** *P* < 0.01; ns, not significant; Wilcoxon rank sum test. **f.** Track views showing RNA-seq signals in *Chd4* Gctrl and *Chd4* GcKO oocytes at P10, on *Stra8* and *Bid* loci. The y-axis represents normalized tag counts for bulk RNA-seq in each sample. Data ranges are shown in brackets.

To determine the function of CHD4 in maintenance of ovarian reserve and survival of GOs, NGOs and GOs were isolated from P10 ovaries, and RNA-seq analysis was performed (Supplementary Fig. 3c). In the P10 *Chd4* GcKO NGOs, 947 genes were up-regulated, and 744 genes were downregulated (Fig. 6d, left, Supplementary Data 4). In the P10 *Chd4* GcKO GOs, 1,938 genes were up-regulated, and 1,420 genes were down-regulated (Fig. 6d, right, Supplementary Data 4). Gene ontology enrichment analyses show that up-regulated genes in P10 *Chd4* GcKO NGOs were associated with apoptotic cell clearance (Supplementary Fig. 3d). In addition, up-regulated genes in P10 *Chd4* GcKO GOs were enriched with genes involved in synaptonemal complex assembly, which is related to MPI (Supplementary Fig. 3d). Female MPI-specific genes were up-regulated in *Chd4* GcKO GOs (Fig. 6e). Similar to *Chd4* DcKO, *Stra8* expression was increased in both P10 NGOs and GOs in *Chd4* GcKO (Fig. 6f). We also examined apoptosis-related genes and found that the expression of a pro-apoptotic gene, *Bid* ^43^, a key player in apoptosis, was increased in both NGOs and GOs in *Chd4* GcKO (Fig. 6f). In summary, CHD4 is essential for oocyte survival and maintenance of ovarian reserve by repressing a group of the MPI genes and apoptosis genes.

## CHD4 repressed apoptosis-related genes for male germ cell survival

After determining the function of CHD4 in the female germline, we finally sought to address whether CHD4 has a common function in the female and male germline. In the male germline, around the time of birth, mitotically arrested prospermatogonia resume active cell cycle and transition to spermatogonia after birth, which sustains long-term fertility of males by stem self-renewal^44^. Recent studies using germline-specific conditional knockout of CHD4 revealed that CHD4 is required for the survival of undifferentiated spermatogonia^20, 21^. Consistent with these studies, our *Chd4* DcKO males (Fig. 7a) showed germ cell depletion that became evident at P3 testes (Fig. 7b).

**Fig 7.**
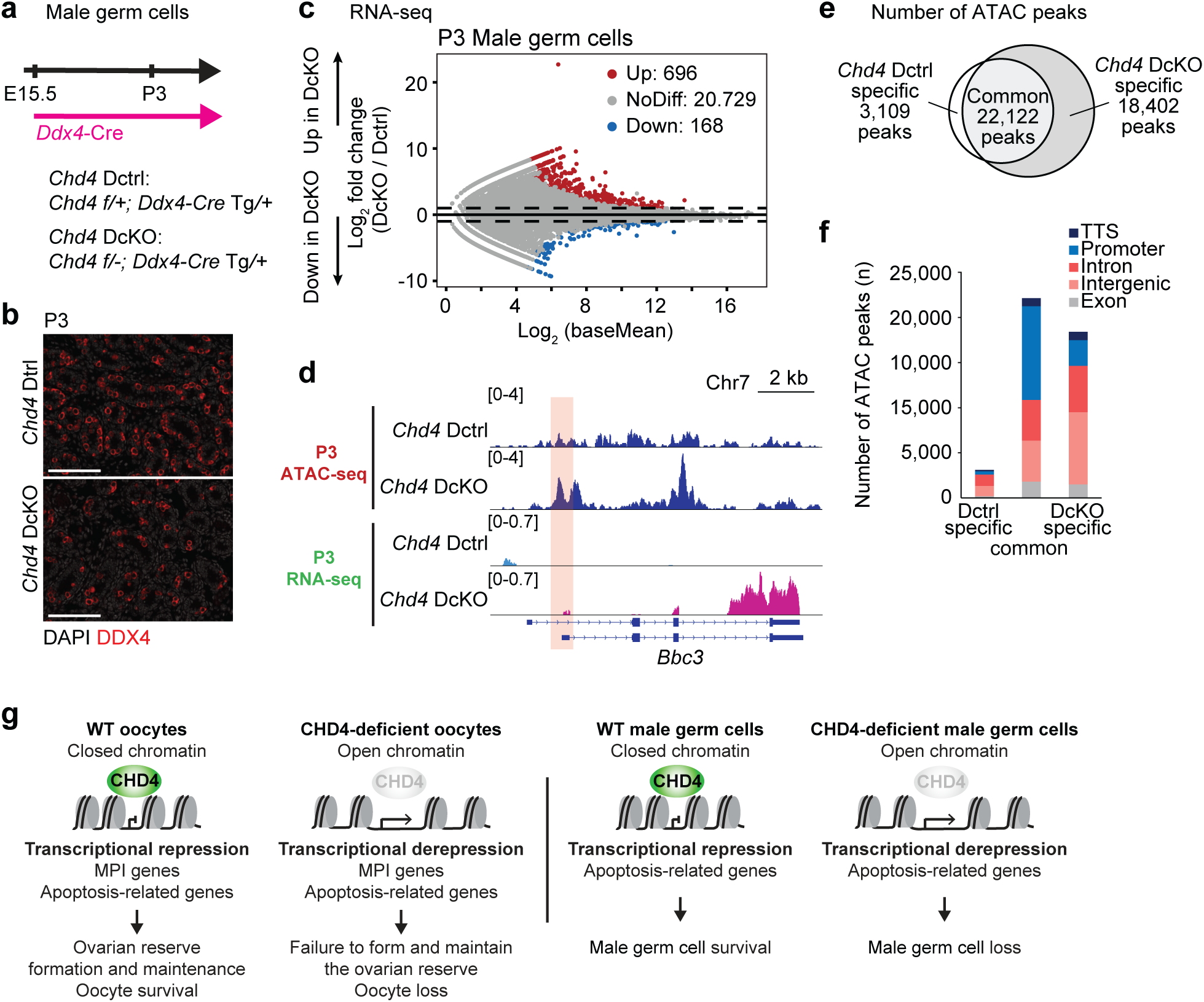
CHD4 suppresses pro-apoptotic genes for male germ cell survival, and summary model. **a.** Schematic for mouse models and experiments. **b.** Immunostaining of DDX4 (red) in testicular sections in *Chd4* Dctrl and *Chd4* DcKO at P3. Bars: 100 μm. Three mice were analyzed for each genotype at each time point, and representative images are shown. **c.** Comparison of transcriptomes between *Chd4* Dctrl and *Chd4* DcKO undifferentiated male germ cells at P3. Two independent biological replicates were examined for RNA-seq. Differentially expressed genes (DEGs: Log2FoldChange > 1, Padj < 0.05, binominal test with Benjamini–Hochberg correction) are colored (red: upregulated in *Chd4* DcKO undifferentiated male germ cells; blue: downregulated in *Chd4* DcKO undifferentiated male germ cells). **d.** Representative track views of *Bbc3* locus in P3 undifferentiated male germ cells of indicated genotypes. Data ranges are shown in brackets. Specific ATAC-seq peak regions in *Chd4* DcKO are highlighted. **e.** Venn diagram indicates overlap of ATAC-seq peaks between *Chd4* Dctrl and *Chd4* DcKO undifferentiated male germ cells at P3. **f.** Numbers and genomic distribution of ATAC-seq peaks in **e**. **g.** Model of CHD4’s function in oocytes and undifferentiated male germ cells.

To examine the genes regulated by CHD4, we isolated undifferentiated male germ cells from P3 testes using a previously established fluorescence-activated cell sorting (FACS) method ^45, 46^ and performed RNA-seq analysis (Supplementary Fig. 4a). In P3 *Chd4* DcKO male germ cells, 696 genes were up-regulated, and 168 genes were down-regulated (Fig. 7c, Supplementary Data 5). Gene ontology enrichment analyses revealed that the genes up-regulated in *Chd4* DcKO male germ cells were associated with “cell morphogenesis” and “regulation of secretion by cell” (Supplementary Fig. 4b). Next, we examined whether the expression of MPI genes is up-regulated by CHD4 deletion, as observed in oocytes, but found no difference in the expression of 104 female MPI genes (Supplementary Fig. 4c). However, a pro-apoptotic gene *Bbc3* and *Gadd45g* were up-regulated in *Chd4* DcKO male germ cells, as was observed in oocytes (Fig. 7d, and Supplementary Fig. 4d). Thus, CHD4 represses the *Bbc3* and *Gadd45* genes in both males and females.

To determine whether CHD4 also represses accessible chromatin in male germ cells, we performed ATAC-seq analysis on P3 *Chd4* DcKO male germ cells (Supplementary Fig. 4e). Chromatin accessibility was increased in *Chd4* DcKO male germ cells compared to P3 *Chd4* Dctrl, as observed in oocytes (Fig. 7e). In addition, P3 *Chd4* DcKO male germ cells -specific ATAC peaks were enriched in intron and intergenic regions, and only slightly in the promoter-TSS region (Fig. 7f). As shown in the track view, chromatin accessibility of the *Bbc3* gene at the TSS was increased (Fig. 7d). These results indicate that CHD4 suppresses expression of apoptosis genes by repressing chromatin accessibility in P3 *Chd4* DcKO male germ cells, leading to cell survival. Taken together, we conclude that CHD4 defines the chromatin state to ensure germ cell survival, enabling the long-term maintenance of female and male germ cells.

## Discussion

The germline must maintain genome integrity to ensure generation of the offspring. Thus, mechanisms underlying long-term maintenance of the germline are critical at sexually dimorphic stages of the germline: one for maintenance of the ovarian reserve in females and another for maintenance of spermatogonial stem cells in males. We report here that CHD4 is a critical regulator for the long-term maintenance of the germline in both males and females. In combination with mouse genetics and epigenomic analyses, our study reveals that CHD4 directly binds and closes accessible chromatin at the distal regulatory elements genome-wide. This mechanism underlies regulation of pro-apoptotic genes in both females and males (Fig. 7g). Notably, the female germline is maintained in the ovarian reserve after MPI and CHD4 is required to close the regulatory elements for MPI genes in females but not in males (Fig. 7g). These results highlight the common and distinct features of chromatin regulation in female and male germlines.

We find that CHD4 is required for both formation and maintenance of ovarian reserve. Because CHD4 has a maintenance function after ovarian reserve formation, it is likely that CHD4 continues to associate with chromatin in MPI-arrested NGOs. The histone H3.3 chaperone HIRA, which continues to replace H3.3 in NGOs is critical to maintain the ovarian reserve^47^, suggesting that the chromatin state of NGOs is not so static. Thus, a chromatin remodeler may be required to maintain a dynamic chromatin environment in NGOs. Furthermore, loss of the DNA damage response (DDR) genes has been implicated in ovarian aging, suggesting a possible function of DDR in the ovarian reserve maintenance^48^. Noteworthy is that CHD4 is known to function in the context of DDR^49, 50^. To further clarify the molecular mechanism for CHD4 in formation and maintenance of the ovarian reserve, the composition of the CHD4-containing chromatin remodeling complex needs to be determined to distinguish its function from other chromatin remodelers whose functions are not known in ovarian reserve formation.

Another critical regulator of POT is PRC1 for MPI exit^7^. Notably, MPI genes repressed by CHD4 (Fig. 3d) are also repressed by PRC1^7^, suggesting possible coordination between CHD4 and PRC1 in MPI exit. In this context, CHD4 closes the accessible chromatin at POT. Consistent with this observation, in embryonic stem cells, CHD4-containing NuRD complexes deacetylate histone H3K27 and recruit PRC2, which often functions with PRC1 to facilitate H3K27me3-mediated repression^51^. On the other hand, CHD4 was also enriched at downregulated genes in *Chd4* DcKO NGOs at P1 and P5 (Fig. 5e), suggesting the possible function of CHD4 in gene activation. Indeed, CHD4 also functions in gene activation^23^, raising the possibility that the function of CHD4 is context-dependent for both gene repression and activation.

We also investigated the target sites of CHD4 chromatin remodeling. The majority of CHD4 binding sites and the accessible chromatin sites closed by CHD4 are intergenic regions and introns. We examined *de novo* motifs present in cKO-specific ATAC peaks (i.e., sites closed by the action of CHD4 in wild-type) in both females and males (Supplementary Fig. 4f). In females, the PRDM9 motif, which is often a feature of meiotic recombination sites, was highly enriched, consistent with CHD4 facilitating exit from the MPI program. The ZNF11 motif was commonly enriched in both females and males, suggesting a common program between males and females. Intriguingly, the NFYB and POU3F1 motifs, detected in males, become open in late spermatogenesis^52^. Thus, it is tempting to speculate that the regulatory elements used in late spermatogenesis are remodeled by CHD4 at an early stage, which may represent a mechanism for epigenetic priming often observed in the male germline^53^.

Together, our study reveals a chromatin remodeling mechanism underlying regulatory elements required for key developmental transitions in the germline. A next key question is how these specific sites are determined to be regulated by CHD4. Because transcription factors (FIGLA, FOXO3) and several signaling pathways (Notch, TGF-β, JNK, and hypoxia signaling) are implicated in primordial follicle formation, it will be important to understand how these mechanisms intersect with chromatin remodeling to establish the necessary chromatin states for ovarian reserve formation. Furthermore, given the significant role of the RNA regulatory network in primordial follicle formation^54^, the mechanistic relationship between the RNA regulatory network and chromatin-based cellular memory emerges as an important agenda for future investigation.

## Methods

### Animals

Generation of conditionally deficient *Chd4* DcKO mice, *Chd4 f/-; Ddx4-Cre Tg/+,* were generated from *Chd4 f/f* female crossed with *Chd4 f/+; Ddx4-Cre Tg/+* males, and *Chd4* Dctrl mice used in experiments were *Chd4 f/+; Ddx4-Cre Tg/+* littermate. Generation of conditionally deficient *Chd4* GcKO mice, *Chd4 f/f; Gdf9-iCre Tg/+,* were generated from *Chd4 f/f* female crossed with *Chd4 f/f; Gdf9-iCre Tg/+* males, and *Chd4* Gctrl mice used in experiments were *Chd4 f/f* littermate. Generation of *Chd4* floxed alleles (*Chd4 f/f*) were reported previously^23^. Mice were maintained on a mixed genetic background of C57BL/6 and DBA2. Ddx4-Cre transgenic mice were purchased from the Jackson Laboratory^24^. For ATAC-seq and CUT&Tag, *Chd4 f/f; Stella*-GFP *Tg/+* mice were generated from *Chd4 f/f* mice crossed with *Stella*-GFP *Tg/+* mice. *Stella*-GFP transgenic mice were obtained from Dr. M. Azim Surani^55^. For each experiment, a minimum of three mice was analyzed. Mice were maintained on a 12:12 light: dark cycle in a temperature and humidity-controlled vivarium (22 ± 2 °C; 40–50% humidity) with free access to food and water in the pathogen-free animal care facility. Mice were used according to the guidelines of the Institutional Animal Care and Use Committee (IACUC: protocol no. IACUC 21931 and 23545) at the University of California, Davis.

### Oocyte collection

The P1, P5, or P10 female pups were collected, and ovaries were harvested by carefully removing oviducts and ovarian bursa in PBS. Ovaries were digested in 200 μl TrypLE™ Express Enzyme (1X) (Gibco, 12604013) supplemented with 0.3 mg/ml Collagenase Type 1 (Worthington, CLS-1) and 10 mg/ml DNase I (Sigma, D5025) and incubated at 37°C for 25 min with gentle agitation. After incubation, the ovaries were dissociated by gentle pipetting using the Fisherbrand^TM^ Premium Plus MultiFlex Gel-Loading Tips until no visible tissue pieces. 2 ml DMEM/F-12 medium (Gibco, 11330107) supplemented with 10% FBS (HyClone, SH30396.03) were then added to the suspension to stop enzyme reaction. Cell suspension was seeded onto a 60 mm tissue culture dish (Falcon, 353002). The cells were allowed to settle down for 15 min at 37°C; 5% CO2 in the incubator before being transferred under the microscope (Nikon, SMZ1270). For RNA-seq, based on morphology and diameter, non-growing and growing oocytes were manually picked up, washed in M2 medium (Sigma, M7167), and transferred into the downstream buffer by mouth pipette. For ATAC-seq and CUT&Tag, P1 non-growing oocytes expressing a *Stella*-GFP transgene were collected by FACS (SONY SH800S).

### Histology and Immunostaining

For the preparation of paraffin blocks, ovaries, and testis were fixed with 4% paraformaldehyde overnight at 4 °C. Ovaries and testis were dehydrated and embedded in paraffin. For histological analysis, 5 µm-thick paraffin sections were deparaffinized and stained with hematoxylin (Sigma, MHS16) and eosin (Sigma, 318906). For immunostaining, 5 µm-thick paraffin sections were deparaffinized and autoclaved in target retrieval solution (DAKO) for 10 min at 121 °C. Sections were blocked with Blocking One Histo (Nacalai) for 30 min at room temperature and then incubated with primary antibodies as outlined below: mouse anti-CHD4 (1:500, Abcam, ab70469), rabbit anti-DDX4 (1:500, Abcam, ab13840), goat anti-CD117/c-kit (1:200, R&D, AF1356), rabbit anti-Cleaved Caspase-3 (1:200, Cell Signaling Technology, #9661), rabbit anti-FOXO3 (1:200, Cell Signaling Technology, #2497) overnight at 4 °C. Sections were washed with PBST (PBS containing 0.1% Tween 20) three times at room temperature for 5 min and then incubated with the corresponding secondary (Invitrogen) at 1:500 dilution for 1 h at room temperature.

Finally, sections were counterstained with DAPI and mounted using 20 μL undiluted ProLong Gold Antifade Mountant (ThermoFisher Scientific, P36930). Images were obtained by an all-in-one fluorescence microscope (BZ-X810, KEYENCE) equipped with an optical sectioning module (BZ-H4XF, KEYENCE).

### Quantification of ovarian follicles

For counting the number of follicles, paraffin-embedded ovaries were serially sectioned at 5 μm thickness, and all sections were mounted on slides. 5 µm-thick paraffin serially sections were deparaffinized and stained with hematoxylin and eosin. Ovarian follicles at different developmental stages, including primordial (type 1 and type 2) as non-growing oocytes, and primary (type 3) and pre-antral (type 4 and type 5) as growing oocytes, were counted in every fifth section of the collected sections from one ovary, based on the standards established method^56^. In each section, only those follicles in which the nucleus of the oocyte was clearly visible were counted, and the cumulative follicle counts were multiplied by a correction factor of 5 to represent the estimated number of follicles in an ovary.

### Meiotic chromosome spreads and immunofluorescence

Chromosome spreads of oocytes from neonatal ovaries were prepared as described^7^. Briefly, ovaries were digested in 200 μl TrypLE™ Express Enzyme (1X) supplemented with 0.3 mg/ml Collagenase Type 1 and 10 mg/ml DNase I and incubated at 37°C for 25 min with gentle agitation. After incubation, the ovaries were dissociated by gentle pipetting using the Fisherbrand^TM^ Premium Plus MultiFlex Gel-Loading Tips until no visible tissue pieces. 2 ml DMEM/F-12 medium supplemented with 10% FBS was added to the suspension to stop enzyme reaction. Cell suspension was incubated in hypotonic extraction buffer [HEB: 30 mM Tris base, 17 mM trisodium citrate, 5 mM ethylenediaminetetraacetic acid (EDTA), 50 mM sucrose, 5 mM dithiothreitol (DTT), 1× cOmplete Protease Inhibitor Cocktail (Sigma, 11836145001), 1× phosphatase inhibitor cocktail 2 (Sigma, P5726-5ML), pH 8.2] on ice for 10 min.

30 µL of the suspension was applied to positively charged slides (Probe On Plus: Thermo Fisher Scientific, 22-230-900); before application of the suspension, the slides had been incubated in chilled fixation solution (2% paraformaldehyde, 0.1% Triton X-100, 0.02% sodium monododecyl sulfate, adjusted to pH 9.2 with sodium borate buffer). The slides were placed in “humid chambers” overnight at room temperature. Then, the slides were washed twice in 0.4% Photo-Flo 200 (Kodak, 146-4510), 2 min per wash. Slides were dried completely at room temperature before staining or storage in slide boxes at −80 °C.

## Flow cytometry and cell sorting

Flow cytometric experiments and cell sorting were performed using SH800S (SONY), with antibody-stained testicular single-cell suspensions prepared as described previously. Data were analyzed using SH800S software (SONY) and FCS Express 7 (De Novo Software).

For ATAC-seq and CUT&Tag, P1 oocytes were collected using the *Stella*-GFP transgene. To prepare single cells suspension for cell sorting, ovaries were digested in 200 μl TrypLE™ Express Enzyme (1X) supplemented with 0.3 mg/ml Collagenase Type 1 and 10 mg/ml DNase I and incubated at 37°C for 25 min with gentle agitation. After incubation, the ovaries were dissociated by gentle pipetting using the FisherbrandTM Premium Plus MultiFlex Gel-Loading Tips until no visible tissue pieces. 2 ml DMEM/F-12 medium supplemented with 10% FBS was added to the suspension to stop enzyme reaction. Cells were suspended in FACS buffer (PBS containing 2% FBS) and filtered into a 5 ml FACS tube through a 35 μm nylon mesh cap (Falcon, 352235). GFP^+^ oocytes were collected after removing small and large debris in FSC-A versus SSC-A gating and doublets in FSC-W versus FSC-H gating.

Collection of male germ cells was modified from described^57^. Briefly, to prepare single cells suspension for cell sorting, detangled seminiferous tubules from P3 mouse testes were incubated in 1× Krebs-Ringer Bicarbonate Buffer (Sigma, K4002) supplemented with 1.5 mg/ml Collagenase Type 1 and 0.04 mg/ml DNase I at 37°C for 15 min with gentle agitation and dissociated using vigorous pipetting. Then add 0.75mg/ml Hyaluronidase (Sigma, H3506) and incubate at 37°C for 10 min with gentle agitation and dissociated using vigorous pipetting.10 ml DMEM/F-12 medium supplemented with 10% FBS was added to the suspension to stop enzyme reaction. The cell suspension was washed with 10 ml FACS buffer three times by centrifugation at 300 × g for 5 min and filtered through a 70 μm nylon cell strainer (Falcon, 352350). The cell suspension was stained with cocktails of antibodies diluted with FACS buffer listed as follows: PE-conjugated anti-mouse/human CD324 (E-Cadherin) antibody (1:500, Biolegend, 147303) and FITC-conjugated anti-mouse CD9 antibody (1:500, Biolegend, 124808). After 50min incubation on ice, cells were washed with 10 ml FACS buffer three times by centrifugation at 300 × g for 5 min and filtered into a 5 ml FACS tube through a 35 μm nylon mesh cap. 7-AAD Viability Stain (Invitrogen, 00-6993-50) and 0.01 mg/ml DNase I was added to cell suspension for the exclusion of dead cells. Samples were kept on ice until sorting. Cells were analyzed after removing small and large debris in FSC-A versus SSC-A gating, doublets in FSC-W versus FSC-H gating, and 7AAD^+^ dead cells. Then, the desired cell population was collected in gates and determined based on antibody staining.

### RNA-seq library generation and sequencing

RNA-seq libraries of oocytes from P1, P5, and P10 ovaries were prepared as described^7^; briefly, 500 non-growing and 100 growing oocytes isolated from ovaries were pooled as one replicate, and two independent biological replicates were used for RNA-seq library generation. Total RNA was extracted using the RNeasy Plus Micro Kit (QIAGEN, Cat # 74034) according to the manufacturer’s instructions. Library preparation was performed with NEBNext® Single Cell/Low Input RNA Library Prep Kit for Illumina® (NEB, E6420S) according to the manufacturer’s instruction. Prepared RNA-seq libraries were sequenced on the HiSeq X system (Illumina) with paired-ended 150-bp reads.

### ATAC-seq library generation and sequencing

ATAC-seq libraries of germ cells were prepared as described^39^; briefly, 10,000 FACS-sorted cells were isolated from P1 ovaries or P3 testis and pooled as one replicate, and two independent biological replicates were used for ATAC-seq library generation. Samples were lysed in 50 μl of lysis buffer (10 mM Tris–HCl (pH 7.4), 10 mM NaCl, 3 mM MgCl2, and 0.1% NP-40, 0.1% Tween-20, and 0.01% Digitonin) on ice for 10 min. Immediately after lysis, the samples were spun at 500 × g for 10 min at 4 °C and the supernatant removed. The sedimented nuclei were then incubated in 10 μl of transposition mix (0.5 µl homemade Tn5 transposase (∼1μg/μl), 5 µl 2× TD buffer (10 mM Tris–HCl (pH 7.6), 10 mM MgCl2, and 20% Dimethyl Formamide), 3.3 µl PBS, 0.1 μl 1% digitonin, 0.1 μl 10% Tween-20, and 1 μl water) at 37 °C for 30 min in a thermomixer with shaking at 500 rpm. After tagmentation, the transposed DNA was purified with a MinElute kit (Qiagen). Polymerase chain reaction (PCR) was performed to amplify the library using the following conditions: 72 °C for 3 min; 98°C for 30 s; thermocycling at 98 °C for 10 s, 60 °C for 30 s, and 72 °C for 1 min. qPCR was used to estimate the number of additional cycles needed to generate products at 25% saturation. Seven to eight additional PCR cycles were added to the initial set of five cycles. Amplified DNA was purified by SPRIselect bead (Beckman Coulter). ATAC-seq libraries were sequenced on the HiSeq X ten system (Illumina) with 150-bp paired-end reads.

### CUT&Tag library generation and sequencing

CUT&Tag libraries from P1 oocytes for CHD4 were generated as previously described^41, 58^ (a step-by-step protocol https://www.protocols.io/view/bench-top-cut-amp-tag-kqdg34qdpl25/v3) using CUTANA™ pAG-Tn5 (Epicypher, 15-1017). Briefly, 10,000 FACS-sorted cells were isolated from P1 *Chd4 f/f* ovaries and pooled as one replicate and two independent biological replicates were used for CUT&Tag library generation. The antibodies used were mouse anti-CHD4 (1:50, Abcam, ab70469) and rabbit α-mouse antibody (1:100, Abcam, ab46540). CUT&Tag libraries were sequenced on the NovaSeq X Plus system (Illumina) with 150-bp paired-end reads.

### RNA-seq data processing

Raw paired-end RNA-seq reads after trimming by trimmomatic (version 0.39)^59^ were aligned to the mouse (GRCm38/mm10) genome using by STAR (version STAR_2.5.4b)^60^ with default arguments. All unmapped and non-uniquely mapped reads were filtered out by samtools (version 1.9)^61^ before being subjected to downstream analyses. To quantify aligned reads in RNA-seq, aligned read counts for each gene were generated using featureCounts (v2.0.1), which is part of the Subread package ^62^ based on annotated genes (GENCODE vM25). The TPM values of each gene were for comparative expression analyses and computing the Pearson correlation coefficient between biological replicates using corrplot^63^. To detect differentially expressed genes between CHD4 Dctrl and CHD4 DcKO, or CHD4 Gctrl and CHD4 GcKO, DESeq2 (version 1.42.1)^64^ was used for differential gene expression analyses with cutoffs ≥2-fold change and binominal tests (Padj < 0.05; P values were adjusted for multiple testing using the Benjamini–Hochberg method). Padj values were used to determine significantly dysregulated genes.

To perform GO analyses, we used the online functional annotation clustering tool Metascape^65^ (http://metascape.org). Further analyses were performed with R and visualized as heatmaps using Morpheus (https://software.broadinstitute.org/morpheus, Broad Institute).

### ATAC-seq and CUT&Tag data processing

Raw paired-end ATAC-seq and CUT&Tag reads after trimming by Trim-galore (https://github.com/FelixKrueger/TrimGalore) (version 0.6.7) were aligned to either the mouse (GRCm38/mm10) genomes using bowtie2 (version 2.3.3.1)^66^ with default arguments. The aligned reads were filtered to remove alignments mapped to multiple locations by calling grep with the -v option before being subjected to downstream analyses. PCR duplicates were removed using the ‘MarkDuplicates’ command in Picard tools (version 2.23.8) (https://broadinstitute.github.io/picard/, Broad Institute). To compare replicates, Pearson correlation coefficients were calculated and plotted by ’multiBamSummary bins’ and ’plot correlation’ functions of deepTools (version 3.3.0)^67^. Biological replicates were pooled for visualization and other analyses after validation of reproducibility. Peak calling for ATAC-seq and CUT&Tag data was performed using MACS3 (version 3.0.0a7)^68^ with default arguments. We computed the number of overlapping peaks between peak files using BEDtools^69^ (version 2.28.0) function intersect. To detect genes adjacent to ATAC-seq and CUT&Tag peaks, we used the HOMER (version 4.9.1)^70^ function annotatePeaks.pl. The deeptools^67^ was used to draw tag density plots and heatmaps for reads enrichments. To visualize ATAC-seq and CUT&Tag data using the Integrative Genomics Viewer (Broad Institute)^71^, BPM normalized counts data were created from sorted BAM files using the deeptools^67^. To perform functional annotation enrichment of CHD4, we used GREAT tools^72^.

### Statistics

Statistical methods and P values for each plot are listed in the figure legends and/or in the Methods. In brief, all grouped data are represented as mean ± SD. All box-and-whisker plots are represented as center lines (median), box limits (interquartile range; 25th and 75th percentiles), and whiskers (maximum value not exceeding 1.5x the interquartile range (IQR) from the hinge) unless stated otherwise. Statistical significance for pairwise comparisons was determined using two-sided Mann–Whitney U-tests and two-tailed unpaired t-tests. Next-generation sequencing data (RNA-seq, ATAC-seq, and CUT&Tag) were based on two independent replicates. No statistical methods were used to predetermine sample size in these experiments. Experiments were not randomized, and investigators were not blinded to allocation during experiments and outcome assessments.

## Data availability

The raw data of quantifications presented in the main figures and supplementary figures are provided as "Source data files". RNA-seq data reported in this study were deposited to the Gene Expression Omnibus (accession no. GSE273309). Source data are provided with this paper.

## Code availability

Source code for all software and tools used in this study with documentation, examples, and additional information, is available at the URLs listed below.

trimmomatic [http://www.usadellab.org/cms/?page=trimmomatic]

STAR [https://github.com/alexdobin/STAR]

featureCounts [http://subread.sourceforge.net]

DESeq2 [https://bioconductor.org/packages/release/bioc/html/DESeq2.html]

corrplot [https://github.com/taiyun/corrplot]

ggplot2 [https://github.com/tidyverse/ggplot2]

Metascape [http://metascape.org]

Morpheus [https://software.broadinstitute.org/morpheus/]

Trim-galore [https://github.com/FelixKrueger/TrimGalore]

Bowtie2 [https://github.com/BenLangmead/bowtie2]

Picard [https://broadinstitute.github.io/picard/]

deepTools [https://github.com/deeptools/deepTools]

MACS3 [https://github.com/macs3-project/MACS]

Bedtools [https://github.com/arq5x/bedtools2]

HOMER [http://homer.ucsd.edu/homer/index.html]

GREAT [http://great.stanford.edu/public/html/]

## Acknowledgments

We thank Tyler Broering and Hironori Abe for their contribution to the initial stage of this project and members of the Namekawa laboratory for discussion and helpful comments regarding this manuscript. We also thank N. Hunter and T. Yoshida for discussion, Katia Georgopoulos for providing the *Chd4* floxed mice, M. Azim Surani for providing *Stella*-GFP transgenic mice, and S. Maezawa for providing the homemade Tn5 transposase for ATAC-seq. Funding sources: Japan Society for Promotion of Science Overseas Research Fellowships, Lalor Foundation Postdoctoral Fellowship, Global Consortium for Reproductive Longevity and Equality (GCRLE) Postdoctoral Fellowship grant number 2223 to Y. M.; UC Davis startup fund, and NIH R35 GM141085 to S.H.N; NIH R21 HD110146 to S.H.N and R.M.S.

## Author information

### Contributions

Y.M. and S.H.N. designed the study. Y.M., M.H., Y.K., A.L.B., and A.S.F. performed experiments and analyzed the data. Y.M., M.H., Y.K., R.M.S., and S.H.N. interpreted the results. Y.M., R.M.S., and S.H.N wrote the manuscript with critical feedback from M.H., Y.K. R.M.S., and S.H.N. supervised the project.

### Corresponding authors

Correspondence to Satoshi H. Namekawa and Richard M. Schultz

