## Supplementary material for "Chromatin remodeler CHD4 establishes chromatin states required for ovarian reserve formation, maintenance, and germ cell survival": Sup Figures

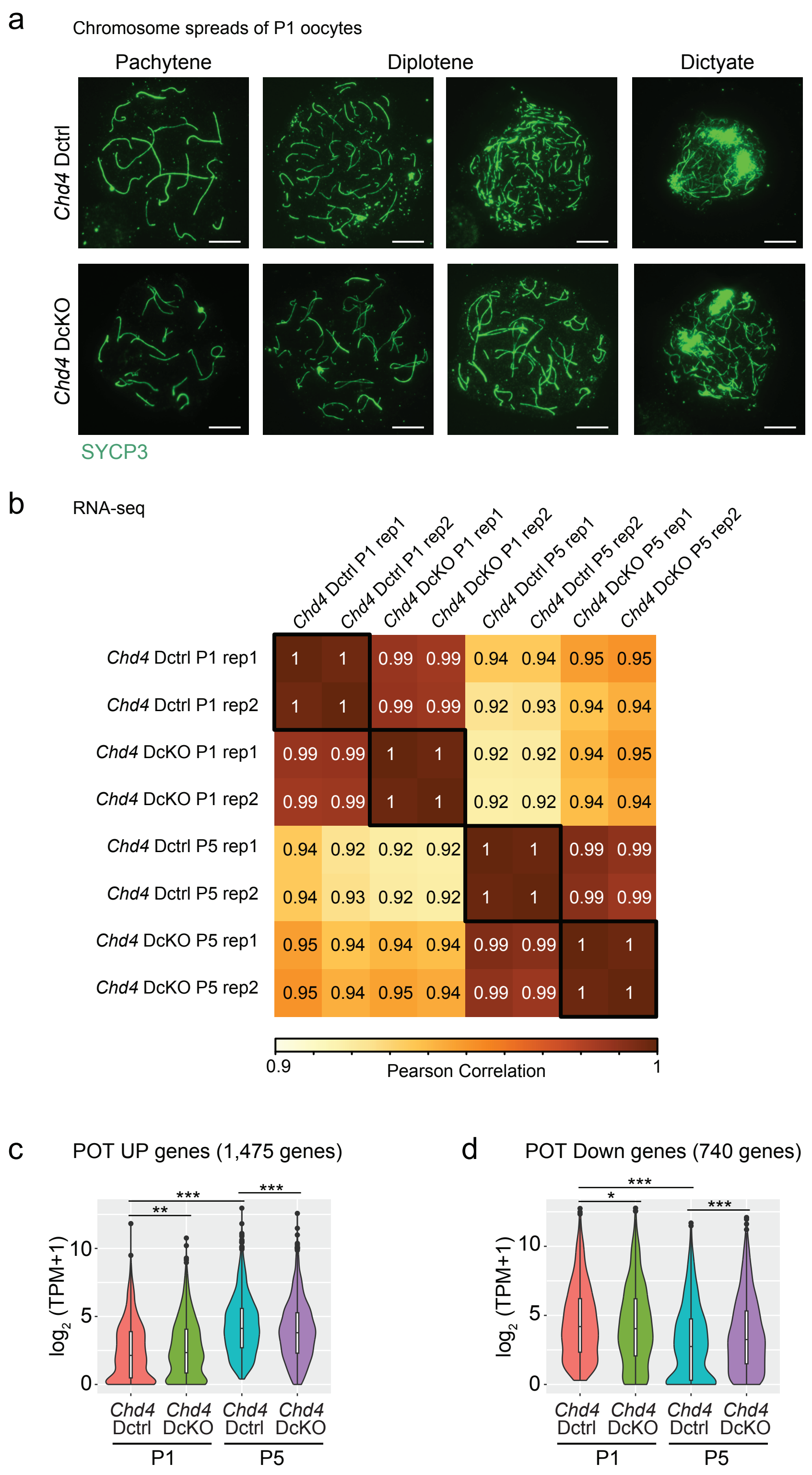
a
Chromosome spreads of P1 oocytes
Pachytene
Diplotene
Dictyate
Chd4 Dctrl
Chd4 DcKO
SYCP3
b
RNA-seq
Chd4 Dctrl P1 rep1
Chd4 Dctrl P1 rep2
Chd4 DcKO P1 rep1
Chd4 DcKO P1 rep2
Chd4 Dctrl P5 rep1
Chd4 Dctrl P5 rep2
Chd4 DcKO P5 rep1
Chd4 DcKO P5 rep2
1 1 0.99 0.99 0.94 0.94 0.95 0.95
1 1 0.99 0.99 0.92 0.93 0.94 0.94
0.99 0.99 1 1 0.92 0.92 0.94 0.95
0.99 0.99 1 1 0.92 0.92 0.94 0.94
0.94 0.92 0.92 0.92 1 1 0.99 0.99
0.94 0.93 0.92 0.92 1 1 0.99 0.99
0.95 0.94 0.94 0.94 0.99 0.99 1 1
0.95 0.94 0.95 0.94 0.99 0.99 1 1
0.9
Pearson Correlation
1
c
POT UP genes (1,475 genes)
log2 (TPM+1)
0
5
10
**
***
***
Chd4 Dctrl
Chd4 DcKO
Chd4 Dctrl
Chd4 DcKO
P1
P5
d
POT Down genes (740 genes)
log2 (TPM+1)
0
5
10
***
*
***
Chd4 Dctrl
Chd4 DcKO
Chd4 Dctrl
Chd4 DcKO
P1
P5

**Supplementary Fig. 1. Analysis of *Chd4* Dctrl and *Chd4* DcKO perinatal oocytes at P1 and P5.**

**a.** Chromosome spreads of P1 oocytes in *Chd4* Dctrl and *Chd4* DcKO immunostained for SYCP3. Bars: 10 μm. Three mice were analyzed for each genotype at each time point, and representative images are shown.

**b.** Heatmap indicates reproducibility between biological replicates in RNA-seq dataset. Pearson correlation values are computed by the "corrplot" package in R.

**c, d.** Violin plots with a Box plot indicate TPM values for POT UP (C, 1,475 genes) and POT Down (D, 740 genes) genes in *Chd4* Dctrl and *Chd4* DcKO oocytes at P1 and P5. POT UP and Down genes were differentially expressed genes (DEGs: Log2FoldChange > 1, Padj < 0.05, binominal test with Benjamini–Hochberg correction) between Ctrl P1 and P5. The central lines represent medians. The upper and lower hinges correspond to the 25th and 75th percentiles. The upper and lower whiskers are extended from the hinge to the largest value no further than the 1.5x inter-quartile range (IQR) from the hinge. *** P < 0.001; ** P < 0.01; * P < 0.05; Wilcoxon rank sum test.

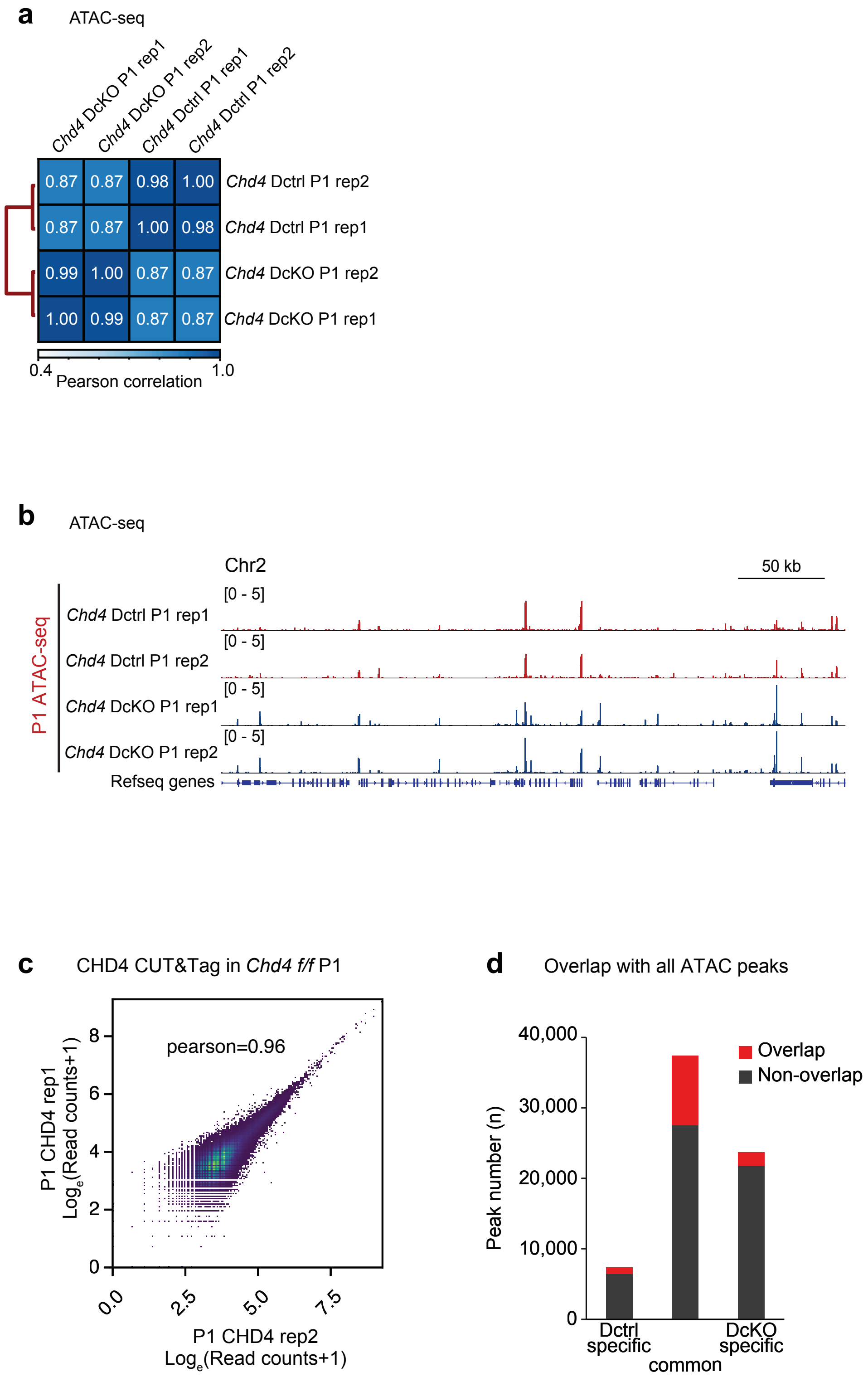
a
ATAC-seq
Chd4 DcKO P1 rep1
Chd4 DcKO P1 rep2
Chd4 Dctrl P1 rep1
Chd4 Dctrl P1 rep2
0.87 0.87 0.98 1.00 Chd4 Dctrl P1 rep2
0.87 0.87 1.00 0.98 Chd4 Dctrl P1 rep1
0.99 1.00 0.87 0.87 Chd4 DcKO P1 rep2
1.00 0.99 0.87 0.87 Chd4 DcKO P1 rep1
0.4
1.0
Pearson correlation
b
ATAC-seq
Chr2
50 kb
P1 ATAC-seq
[0 - 5]
Chd4 Dctrl P1 rep1
[0 - 5]
Chd4 Dctrl P1 rep2
[0 - 5]
Chd4 DcKO P1 rep1
[0 - 5]
Chd4 DcKO P1 rep2
Refseq genes
c
CHD4 CUT&Tag in Chd4 f/f P1
pearson=0.96
P1 CHD4 rep1
Log<sub>e</sub>(Read counts+1)
0
2
4
6
8
0.0
2.5
5.0
7.5
P1 CHD4 rep2
Loge(Read counts+1)
d
Overlap with all ATAC peaks
Overlap
Non-overlap
Peak number (n)
0
10,000
20,000
30,000
40,000
Dctrl specific
common
DcKO specific

**Supplementary Fig. 2. ATACseq and CUT&Tag analysis in *Chd4* Dctrl and *Chd4* DcKO oocytes at P1.**
**a.** Heatmap indicates reproducibility between biological replicates in ATAC-seq dataset. Pearson correlation values are computed by deepTools package.
**b.** Representative track view of ATAC-seq enrichments in biological replicates. The y-axis represents the normalized BPM for ATAC-seq in each sample.
**c.** Scatter plot shows the reproducibility between biological replicates in CUT&Tag data. Pearson correlation values (R) are shown.
**d.** Overlap between all ATAC peaks and CHD4 CUT&Tag peaks within classified ATAC peaks.

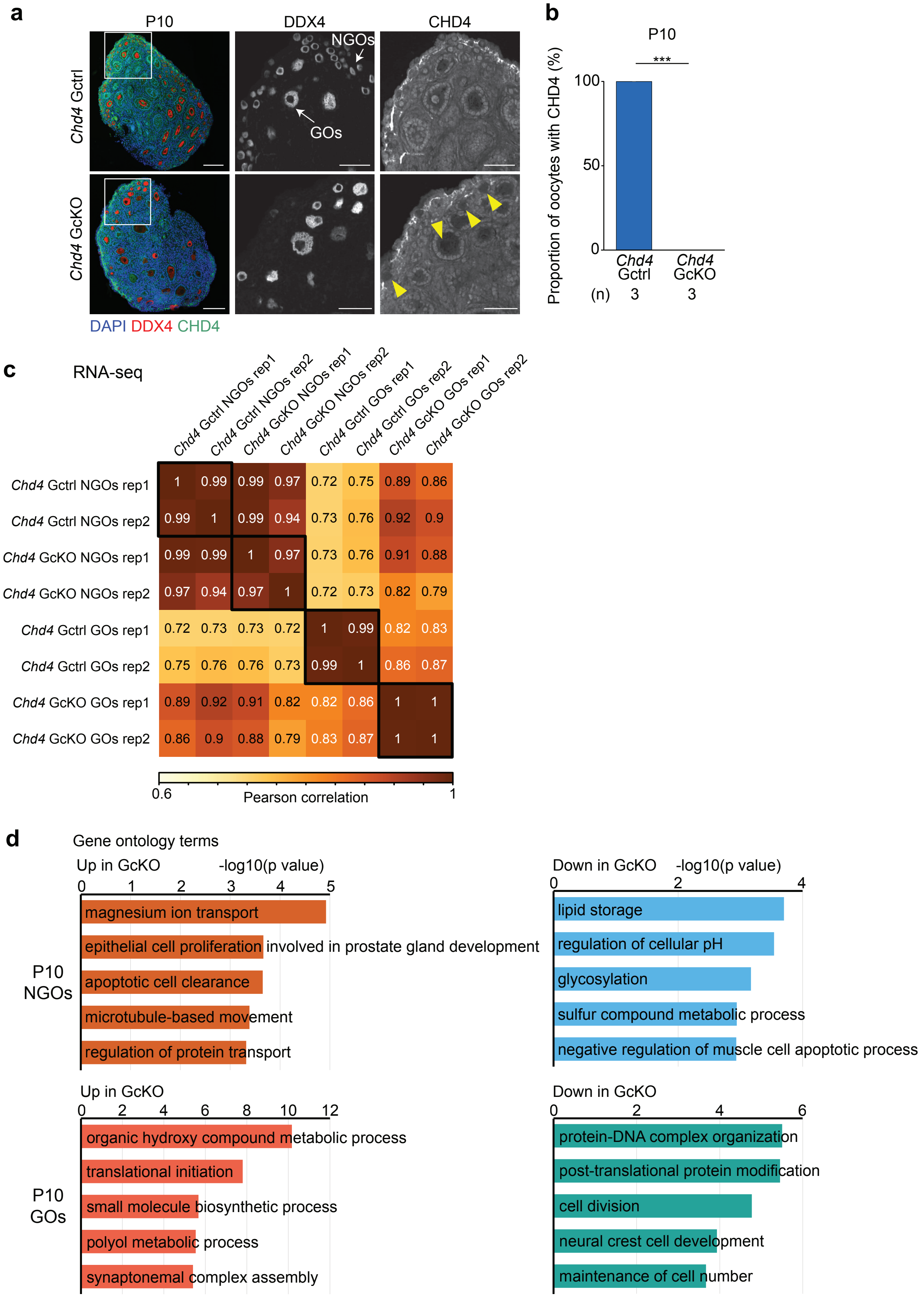


| | Chd4 Gctrl NGOs rep1 | Chd4 Gctrl NGOs rep2 | Chd4 GcKO NGOs rep1 | Chd4 GcKO NGOs rep2 | Chd4 Gctrl GOs rep1 | Chd4 Gctrl GOs rep2 | Chd4 GcKO GOs rep1 | Chd4 GcKO GOs rep2 |
|---|---|---|---|---|---|---|---|---|
| Chd4 Gctrl NGOs rep1 | 1 | 0.99 | 0.99 | 0.97 | 0.72 | 0.75 | 0.89 | 0.86 |
| Chd4 Gctrl NGOs rep2 | 0.99 | 1 | 0.99 | 0.94 | 0.73 | 0.76 | 0.92 | 0.9 |
| Chd4 GcKO NGOs rep1 | 0.99 | 0.99 | 1 | 0.97 | 0.73 | 0.76 | 0.91 | 0.88 |
| Chd4 GcKO NGOs rep2 | 0.97 | 0.94 | 0.97 | 1 | 0.72 | 0.73 | 0.82 | 0.79 |
| Chd4 Gctrl GOs rep1 | 0.72 | 0.73 | 0.73 | 0.72 | 1 | 0.99 | 0.82 | 0.83 |
| Chd4 Gctrl GOs rep2 | 0.75 | 0.76 | 0.76 | 0.73 | 0.99 | 1 | 0.86 | 0.87 |
| Chd4 GcKO GOs rep1 | 0.89 | 0.92 | 0.91 | 0.82 | 0.82 | 0.86 | 1 | 1 |
| Chd4 GcKO GOs rep2 | 0.86 | 0.9 | 0.88 | 0.79 | 0.83 | 0.87 | 1 | 1 |

**Supplementary Fig. 3. Validation and transcriptome analysis of *Gdf9*-iCre in oocytes at P10.**

**a.** Immunostaining of DDX4 (red) and CHD4 (green) in ovaries of *Chd4* Gctrl and *Chd4* GcKO at P10. CHD4 is present only in somatic cells in *Chd4* GcKO. Bars: 100 μm (50 μm in the boxed area). Yellow arrowheads indicate oocytes with no CHD4 expression.

**b.** Quantitative analysis of immunostaining.

**c.** Heatmap indicates reproducibility between biological replicates in RNA-seq dataset. Pearson correlation values are computed by the "corrplot" package in R.

**d.** Gene ontology term enrichments analysis of differentially expressed genes detected in Fig 7d.

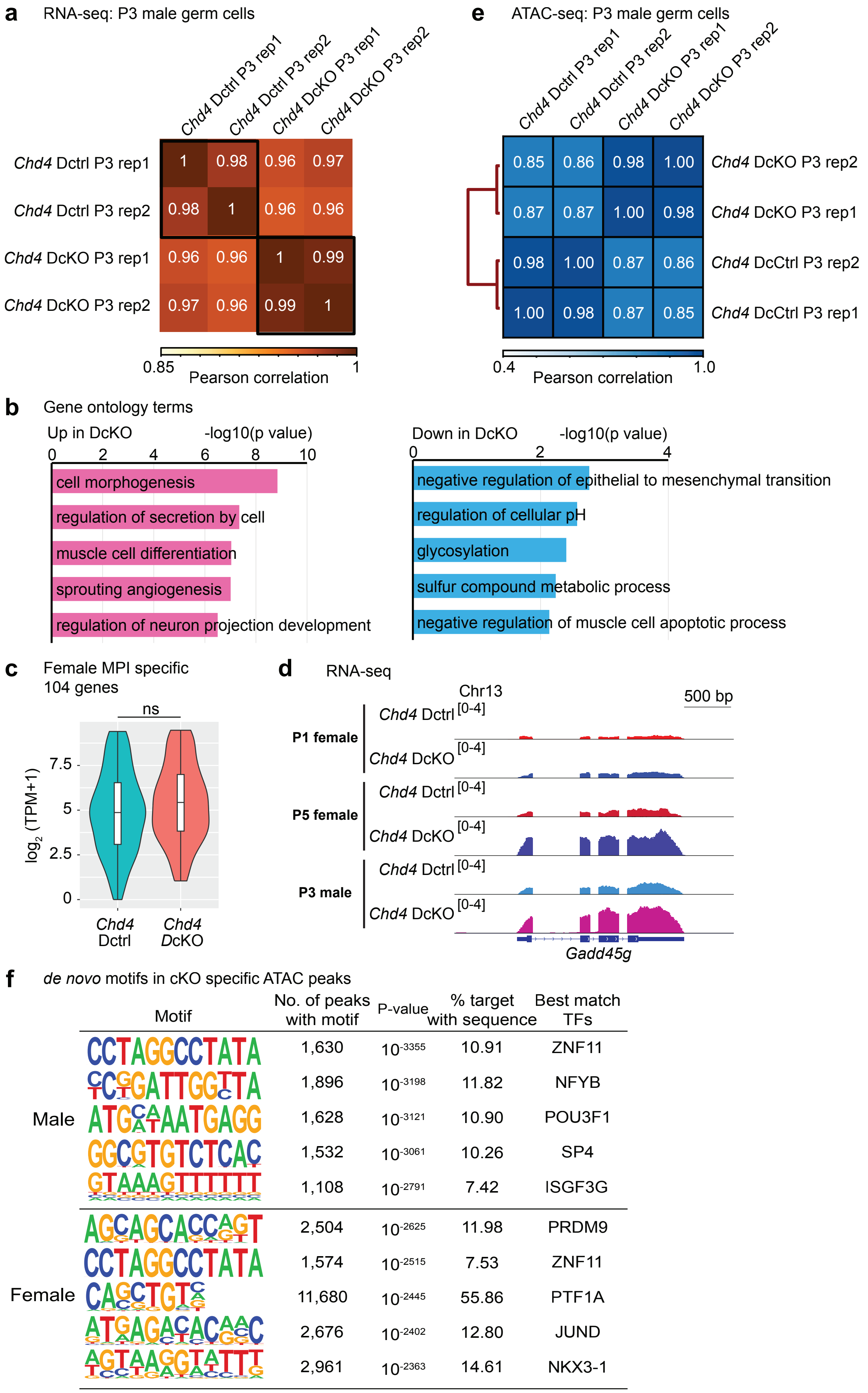
a
RNA-seq: P3 male germ cells
Chd4 Dctrl P3 rep1
Chd4 Dctrl P3 rep2
Chd4 DcKO P3 rep1
Chd4 DcKO P3 rep2
1 0.98 0.96 0.97
0.98 1 0.96 0.96
0.96 0.96 1 0.99
0.97 0.96 0.99 1
0.85
1
Pearson correlation
e
ATAC-seq: P3 male germ cells
Chd4 Dctrl P3 rep1
Chd4 Dctrl P3 rep2
Chd4 DcKO P3 rep1
Chd4 DcKO P3 rep2
0.85 0.86 0.98 1.00 Chd4 DcKO P3 rep2
0.87 0.87 1.00 0.98 Chd4 DcKO P3 rep1
0.98 1.00 0.87 0.86 Chd4 DcCtrl P3 rep2
1.00 0.98 0.87 0.85 Chd4 DcCtrl P3 rep1
0.4
1.0
Pearson correlation
b
Gene ontology terms
Up in DcKO
-log10(p value)
0 2 4 6 8 10
cell morphogenesis
regulation of secretion by cell
muscle cell differentiation
sprouting angiogenesis
regulation of neuron projection development
Down in DcKO
-log10(p value)
0 2 4
negative regulation of epithelial to mesenchymal transition
regulation of cellular pH
glycosylation
sulfur compound metabolic process
negative regulation of muscle cell apoptotic process
c
Female MPI specific
104 genes
ns
7.5
5
2.5
0
log2 (TPM+1)
Chd4 Dctrl
Chd4 DcKO
d
RNA-seq
Chr13
500 bp
P1 female
Chd4 Dctrl [0-4]
Chd4 DcKO [0-4]
P5 female
Chd4 Dctrl [0-4]
Chd4 DcKO [0-4]
P3 male
Chd4 Dctrl [0-4]
Chd4 DcKO [0-4]
Gadd45g
f
de novo motifs in cKO specific ATAC peaks
Motif
No. of peaks with motif
P-value
% target with sequence
Best match TFs
Male
CCTAGGCCTATA 1,630 10-3355 10.91 ZNF11
1,896 10-3198 11.82 NFYB
1,628 10-3121 10.90 POU3F1
1,532 10-3061 10.26 SP4
1,108 10-2791 7.42 ISGF3G
Female
2,504 10-2625 11.98 PRDM9
CCTAGGCCTATA 1,574 10-2515 7.53 ZNF11
11,680 10-2445 55.86 PTF1A
2,676 10-2402 12.80 JUND
2,961 10-2363 14.61 NKX3-1

**Supplementary Fig. 4. RNA- and ATAC-seq analysis in *Chd4* Dctrl and *Chd4* DcKO male germ cell at P3.**
**a.** Heatmap indicates reproducibility between biological replicates in RNA-seq dataset. Pearson correlation values are computed by the "corrplot” package in R.
**b.** Gene ontology term enrichment analysis of differentially expressed genes detected in Fig 7c.
**c.** Violin plots with a Box plot indicate TPM values for female MPI-specific genes (104 genes) in *Chd4* Dctrl and *Chd4* DcKO male germ cells at P3. The central lines represent medians. The upper and lower hinges correspond to the 25th and 75th percentiles. The upper and lower whiskers are extended from the hinge to the largest value no further than the 1.5x inter-quartile range (IQR) from the hinge. ns, not significant; Wilcoxon rank sum test.
**d.** Representative track views of the *Gadd45g* gene locus in P1 and P5 oocytes and P3 male germ cells of indicated genotypes. Data ranges are shown in brackets.
**e.** Heatmap indicates reproducibility between biological replicates in the ATAC-seq dataset. Pearson correlation values are computed by the deepTools package.
**f.** HOMER *de novo* motif analyses of specific ATAC-seq peaks in *Chd4* DcKO male germ cell at P3 and oocytes at P1 for putative TF-binding sites.

**Description of supplementary files**

**Supplementary Data 1**

Differentially expressed genes in *Chd4* Dctrl and *Chd4* DcKO oocytes at P1

**Supplementary Data 2**

Differentially expressed genes in *Chd4* Dctrl and *Chd4* DcKO oocytes at P5

**Supplementary Data 3**

Differentially expressed genes in P1 *Chd4* Dctrl oocyte and P5 *Chd4* Dctrl oocytes during POT

**Supplementary Data 4**

Differentially expressed genes in *Chd4* Gctrl and *Chd4* GcKO oocytes at P10

**Supplementary Data 5**

Differentially expressed genes in *Chd4* Dctrl and *Chd4* DcKO undifferentiated male germ cells at P3
